# StCEL-based platform enables detection and sequencing of native sialoglycoRNA

**DOI:** 10.1101/2025.08.06.668864

**Authors:** Guang-Dong Zeng, Yun-Long Wang, Xu-Hui Chen, Yi Ma, Zhu Zhu, Huan-Lei Liu, Li-Yuan Fan, Hao Nan, Ziwaregul Nur, Min Wei, Xin-Yuan Liu, Li Yi, Jiu-Yu Jin, Yu-Song Cao, Meng-Bo Qiu, Wei-Peng Sun, Yang Bai, Quan-Bo Zhou, Guo-Qing Zhang, Jing-Feng Chen, Jie Shi, Jian Zheng, Yixuan Xie, Peng Li, Xiang-Bo Wan, Xin-Juan Fan

## Abstract

Glycosylated RNAs (glycoRNAs) are an emerging class of biomolecules characterized by glycans attached to RNA molecules. Currently, glycoRNA detection remains largely confined to cellular models, substantially impeding its functional characterization and clinical translation. Here, we developed sialyltransferase-mediated chemical enzymatic labeling (StCEL), in which azido-modified sialic acid was transferred to the natural sialic acid on sialoglycoRNA, enabling the detection of native sialoglycoRNA across diverse biological samples with high specificity and sensitivity. Using StCEL, we found that sialoglycoRNA was significantly upregulated in multiple cancer types. StCEL combined with a microplate reader was applied to detect sialoglycoRNA in large-scale samples, revealing that elevated sialoglycoRNA levels correlate positively with tumor malignancy and poor patient prognosis. Furthermore, StCEL-based sialoglycoRNA enrichment and sequencing identified dysregulated sialoglycoRNAs in tumor tissues. Collectively, StCEL establishes a multifunctional platform that enables sensitive and specific detection of sialoglycoRNA, alongside sequencing analysis, demonstrating significant potential for clinical translation.

## Introduction

Glycosylation is a fundamental cellular mechanism critical to physiological and pathological processes^1–5^. While the functions of glycoprotein, glycolipid, and proteoglycan are well-characterized^6–9^, glycosylation’s potential roles in RNA biology remained unexplored until 2021, when Flynn et al. first identified glycosylated small RNAs (glycoRNAs)^10^. Specifically, these glycoRNAs were sialylated (sialoglycoRNAs) and localized to the cell surface^10^. Subsequent work revealed that 3-(3-amino-3-carboxypropyl) uridine (acp^3^U) was the attachment site for N-glycans^11^. Functionally, Zhang et al. demonstrated that glycoRNAs were critical for neutrophil recruitment to inflammatory sites by promoting neutrophil adhesion and migration through endothelial cells^12^. Moreover, Perr et al. found that glycoRNAs co-assembled with cell surface-localizing RNA-binding proteins (RBPs) to act as specialized molecular gateways for the cellular internalization of cell-penetrating peptides, such as the HIV trans-activator of transcription protein^13^. Nevertheless, understanding of glycoRNA functions and associated molecular mechanisms remains limited, largely attributable to constraints in current glycoRNA detection methods.

Several sialoglycoRNA detection methods have been developed. For sialoglycoRNA imaging, both the sialic acid aptamer-based RNA proximity ligation assay (ARPLA) and the second-generation hierarchical coding strategy (HieCo2) leverage proximity ligation to label sialoglycoRNAs, enabling spatial imaging of single sialoglycoRNA species in situ^14, 15^. Additionally, dual-recognition Förster resonance energy transfer (drFRET) facilitated the imaging of single sialoglycoRNA species in small extracellular vesicles (sEVs)^16^. However, these imaging methods suffer from procedurally cumbersome workflows with quantitatively limited sensitivity, while their individual RNA species detection inherently precludes comprehensive assessment of the global sialoglycoRNA landscape. As for detecting sialoglycoRNA by blotting, Ac_4_ManNAz-based metabolic labeling is the common approach used in glycoRNA discovery; its application is restricted to analyses in living cells or animal models, and thus it is unsuitable for dissected tissues or body fluid samples^10^. Additionally, variations in the incorporation rate of Ac_4_ManNAz limit its application in quantification analysis^17, 18^. Moreover, periodate oxidation and aldehyde ligation (rPAL) oxidizes vicinal diols in sialic acid residues, allowing for the detection of native sialoglycoRNAs, while this approach generates substantial nonspecific RNA labeling by oxidizing the 2′,3′-vicinal diol in RNA’s 3′-terminal ribose^11^. Overall, despite the availability of multiple detection methods, only rPAL enables panoramic profiling of native sialoglycoRNAs. However, rPAL generates substantial nonspecific signal, interfering with sialoglycoRNA analysis and critically impairing subsequent enrichment, thereby precluding its application in sialoglycoRNA sequencing, which is essential to investigate sialoglycoRNA functions and pioneer their applications. Therefore, a metabolic labeling-independent method for native sialoglycoRNA detection in natural samples, especially to enable sialoglycoRNA-seq profiling, is urgently needed.

Here, we developed sialyltransferase-mediated chemical enzymatic labeling (StCEL), a highly sensitive, specific, and innovative method to label native sialoglycoRNA across diverse biological samples. Methodologically, CstⅡ, a *Campylobacter jejuni-*derived α-2,8-sialyltransferase^19–21^, catalyzes CMP-Neu5Az transfer to α-2,3-or α-2,6-linked sialic acid residues^22, 23^, followed by click chemistry-mediated biotin or fluorescent dye labeling to enable the expression level analysis and enrichment of the labeled sialoglycoRNA for RNA-seq. Using StCEL, we discovered that sialoglycoRNA levels were considerably elevated in various cancer types and high sialoglycoRNA expression was associated with poor prognosis in patients with rectal cancer. StCEL-based enrichment coupled with high-throughput sequencing revealed dysregulated sialoglycoRNAs in rectal tumor tissues, implicating sialoglycoRNA involvement in carcinogenesis. These results demonstrate StCEL as a highly sensitive and specific method to measure sialoglycoRNA abundance and to perform sialoglycoRNA-seq in clinical samples, thereby providing a platform to explore sialoglycoRNAs’ physiological and pathological functions.

## Results

### Development of StCEL for detecting native sialoglycoRNAs

To label native sialoglycoRNA for abundance analysis and RNA-seq, we developed a method called StCEL. This method employed CstⅡ, an α-2,8-monosialyltransferase from *Campylobacter jejuni*^19–21^, to specifically transfer azido-modified sialic acid to α-2,3-or α-2,6-linked sialic acid on native sialoglycoRNA^22, 23^, which preserves the original glycosylation and sialylation information (Fig. 1). As expected, StCEL successfully detected sialoglycoRNA signal in HCT 116 and HeLa cells under the conditions previously applied in glycolipid and glycoprotein biosynthesis and detection^23–25^ (Fig. 2a and Extended Data Fig. 1a).

**Fig. 1.**
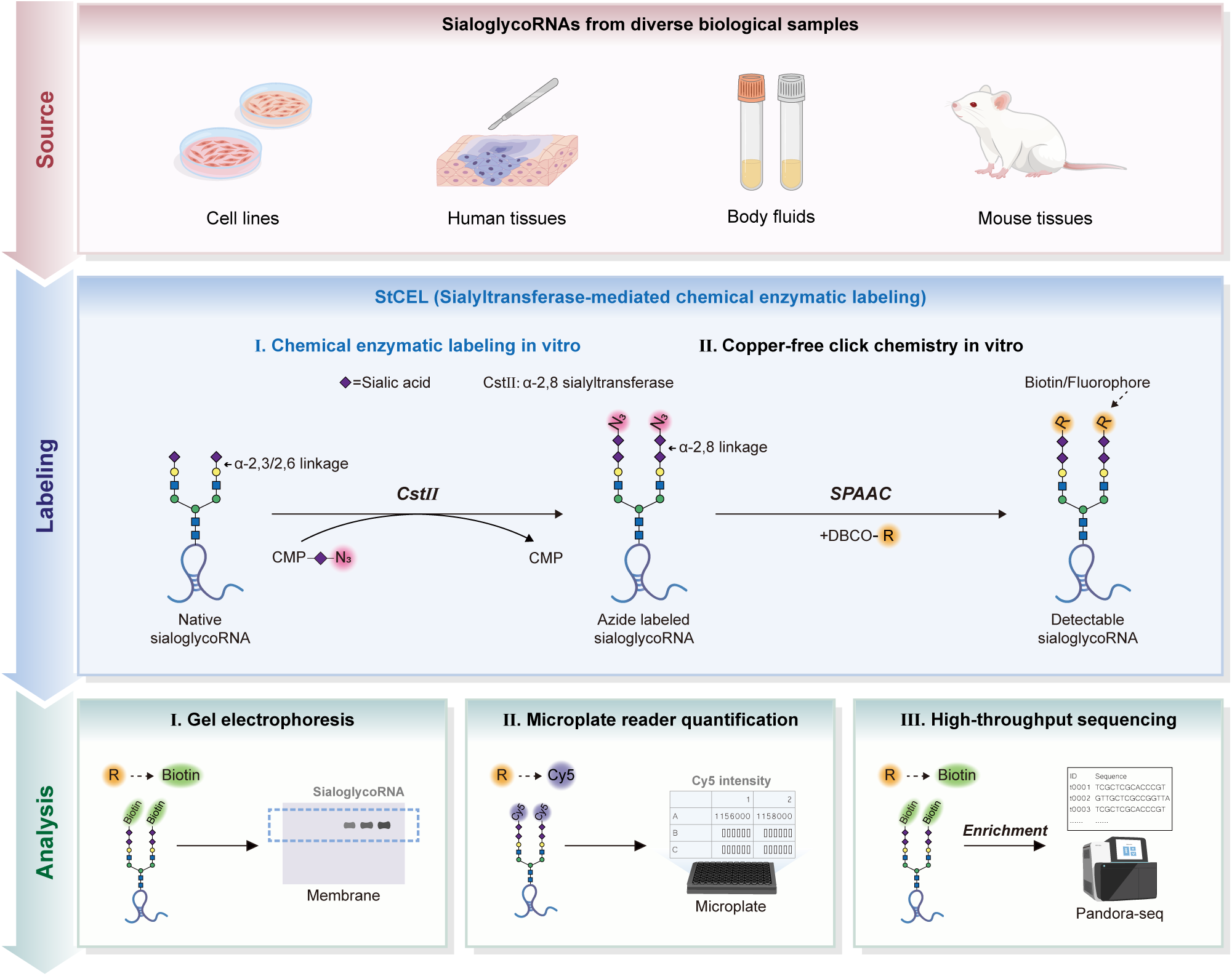
Schematic illustration of the StCEL platform for sialoglycoRNA labeling and analysis. The StCEL platform enables comprehensive analysis of sialoglycoRNA across diverse biological samples. The platform first employs sialyltransferase CstⅡ to catalyze the transfer of azido-modified sialic acid to endogenous sialoglycoRNA, followed by SPAAC labeling with DBCO-biotin or DBCO-fluorophores (e.g., Cy5). This platform supports two functional modes: (1) Detection: Northern blot visualization of biotin/Cy5 tags or high-throughput Cy5 quantification using microplate readers; (2) Enrichment: Coupling to RNA-seq for sequence analysis, thereby enabling comprehensive characterization of sialoglycoRNA.

**Fig. 2.**
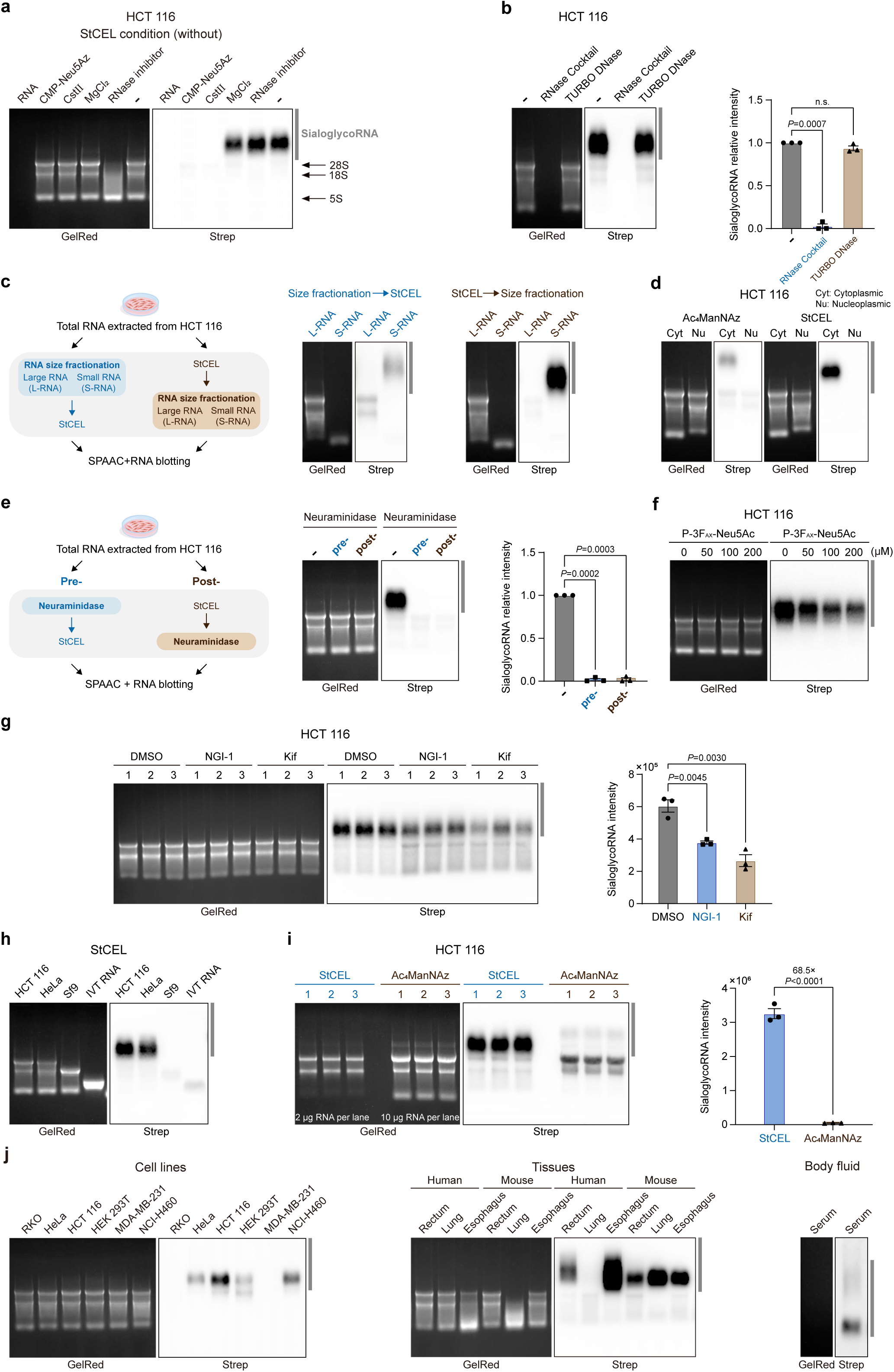
StCEL labels sialoglycoRNA with high specificity and sensitivity. **a**, RNA blot analysis of HCT 116 total RNA labeled with StCEL in the absence of individual component (RNA, CMP-Neu5Az, CstⅡ, MgCl_2,_ or RNase inhibitor). Left panel: total RNA detected by GelRed (in-gel). Right panel: biotinylated sialoglycoRNA detected by HRP-streptavidin (on-membrane). The region of sialoglycoRNA is marked by dark gray vertical bars. **b**, RNA blotting of StCEL-labeled HCT 116 RNA treated with RNase cocktail (A/T1) or TURBO DNase in vitro (left). Each datapoint (biological triplicate) was displayed with the mean ±standard error of the mean (SEM, right), and statistical analysis was performed with paired *t* tests. n.s. means not significant. **c**, Schematic of RNA size fractionation assay before or after StCEL (left). RNA blotting of HCT 116 L-RNA and S-RNA populations labeled with StCEL (right). L-RNA, large RNA (>200 nt); S-RNA, small RNA (<200 nt). **d**, RNA blotting of RNA extracted from cytoplasmic (Cyt) and nucleoplasm (Nu) fractions of HCT 116 cells labeled with Ac_4_ManNAz or StCEL. **e**, Schematic of total RNA treated by α2-3,6,8,9 Neuraminidase A before or after StCEL (left). RNA blotting of StCEL-labeled HCT 116 total RNA treated by α2-3,6,8,9 Neuraminidase A before or after StCEL (middle). Each datapoint (biological triplicate) was displayed with the mean ±SEM (right), and statistical analysis was performed with paired *t* tests. **f**, HCT 116 cells were treated with the indicated concentrations of P-3F_AX_-Neu5Ac for 24 hours before StCEL labeling and RNA blotting. **g**, RNA blotting of StCEL-labeled total RNA from HCT 116 cells treated with N-glycan inhibitors. 0.05% DMSO, 10 μM NGI-1, and 4 μM kifunensine (kif) were used (left). Right panel: each datapoint (biological triplicate) was displayed with the mean ±SEM (right), and statistical analysis was performed with unpaired *t* tests. **h**, RNA blotting of total RNA derived from HCT 116, HeLa, Sf9, and in vitro transcription (IVT) labeled with StCEL. **i**, RNA blotting of total RNA from HCT 116 cells labeled with StCEL or Ac_4_ManNAz (left). Each datapoint (biological triplicate) is displayed with the mean ±SEM (right), and statistical analysis was performed with an unpaired *t*-test. **j**, StCEL-mediated sialoglycoRNA detection in cell lines, human clinical tissues, mouse tissues, and human body fluid.

Next, we assessed the specificity and sensitivity of StCEL. As shown, RNase cocktail treatment effectively abolished the StCEL signal, whereas DNase treatment had no effect (Fig. 2b and Extended Data Fig. 1b). Consistent with the result of previous studies^10, 11^, the StCEL signals were predominantly detected in the small RNA (Fig. 2c and Extended Data Fig. 1c) and cytoplasmic fractions (Fig. 2d and Extended Data Fig. 1d). Furthermore, treatment of RNA with α2-3,6,8,9 Neuraminidase A completely abolished the StCEL signal (Fig. 2e and Extended Data Fig. 1e). Similarly, inhibition of sialic acid modification using P-3F_AX_-Neu5Ac led to a dose-dependent reduction in the StCEL signals (Fig. 2f and Extended Data Fig. 1f). Furthermore, cells treated with N-glycosylation inhibitors (NGI-1 and kifunensine [kif]) significantly attenuated the StCEL signals (Fig. 2g). The StCEL signals was exclusively detected in mammalian cells but absent from both insect Sf9 cells and in vitro-transcribed RNA due to their lack of sialic acid biosynthesis pathways^26^ (Fig. 2h and Extended Data Fig. 1g). Significantly, StCEL showed higher sensitivity than Ac_4_ManNAz labeling, presenting 68.5-and 116-fold stronger signals in HCT 116 and HeLa cells, respectively (Fig. 2i and Extended Data Fig. 1h). Since StCEL labels sialoglycoRNAs post total RNA extraction, it confers broad applicability across diverse biological sources, including cell lines, human clinical tissues, liquid biopsies, and animal models (Fig. 2j). These findings confirm that StCEL can label sialoglycoRNA with high specificity and sensitivity, enabling robust analysis of diverse biological sources.

### StCEL identifies sialoglycoRNAs upregulation in tumor tissues

To demonstrate the broad application scenarios of StCEL, we tested its performance on clinical samples. Consistent with its performance in cellular RNA detection, StCEL showed high specificity in detecting sialoglycoRNA in clinical tissues (Extended Data Fig. 2a-c). Aberrant expression of glycans have been well established as biomarkers for cancer diagnosis and prognosis^27–30^, but the link between sialoglycoRNA expression levels and cancers remains incompletely resolved^14, 16^. Therefore, we first examined the landscape of sialoglycoRNAs’ expression in tumor-normal tissue pairs. Intriguingly, sialoglycoRNAs were upregulated in 80.49% (33/41) of rectal, 72.5% (29/40) of lung, and 70.83% (17/24) of esophageal tumor tissues compared to adjacent normal tissues (Fig. 3a-e; Extended Data Fig. 2d-g and Supplementary Fig. 1a-c). These results suggest that sialoglycoRNA upregulation is a pan-cancer feature. Furthermore, analysis of isolated cells revealed predominant localization of sialoglycoRNA in non-immune (CD45⁻) cells within rectal tumor tissues. (Fig. 3f; Extended Data Fig. 2h and Supplementary Fig. 1d).

**Fig. 3.**
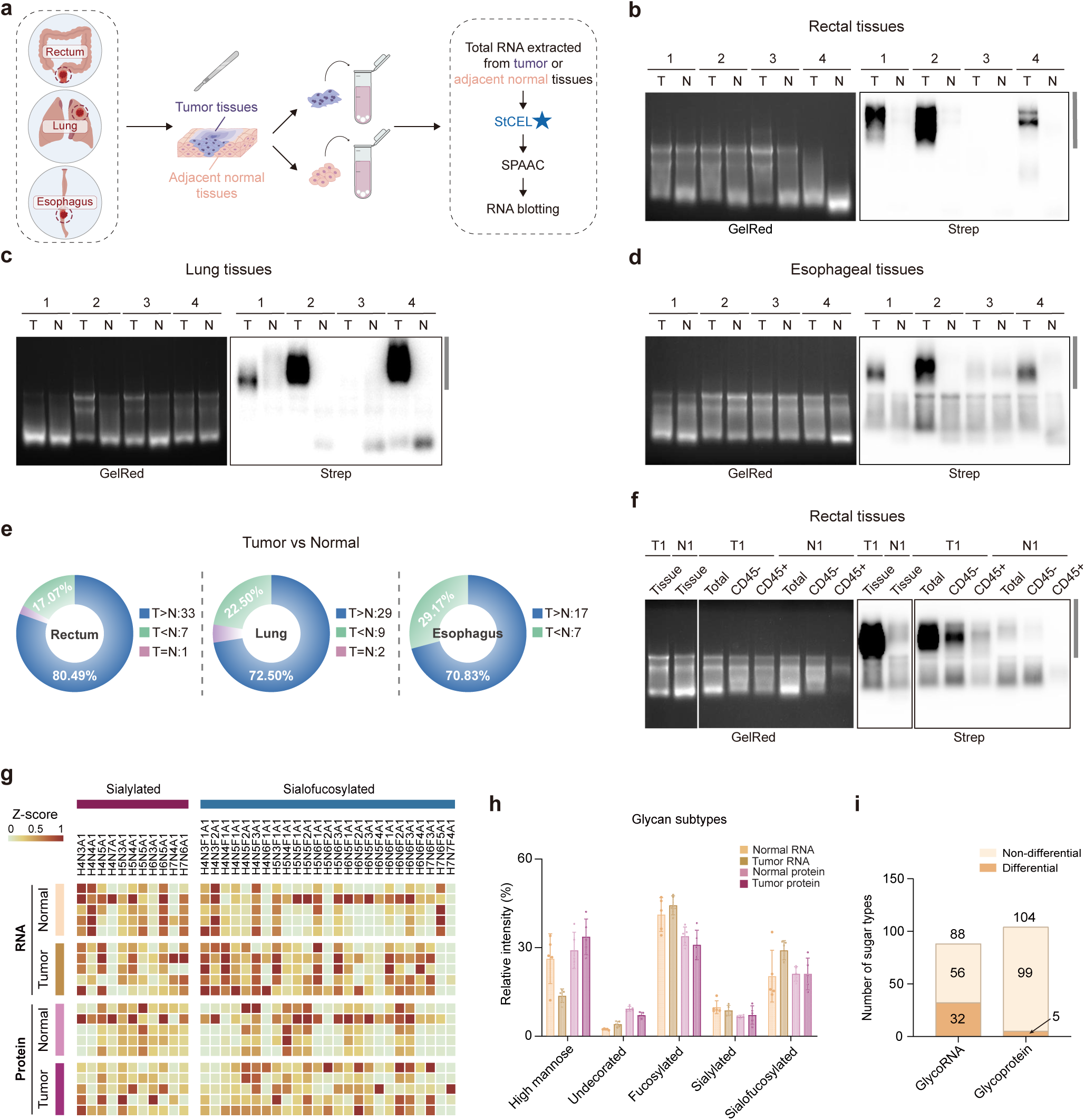
StCEL enables sialoglycoRNA detection in clinical tissues. **a**, Schematic of the sialoglycoRNA detection strategy applied to tumor (T) and paired adjacent normal (N) tissues of rectal, lung, and esophageal cancer patients. The blue pentagram is meant to emphasize the StCEL method. **b,c,d**, Representative RNA blotting of tumor (T) and adjacent normal (N) tissues in rectal cancer (**b**, 4 of 41 pairs shown), lung cancer (**c**, 4 of 40), and esophagus cancer (**d**, 4 of 24). **e**, Pie chart comparing sialoglycoRNA signal distribution between tumor (T) and adjacent normal (N) tissues across cancer cohorts (rectum, n=41; lung, n=40; esophagus, n=24). Data were derived from the grayscale values of the sialoglycoRNA signals in Fig. 3b**-d**, **Extended Data Fig. 2d-f**, and **Supplementary Fig. 1a-c. f**, RNA blotting of StCEL-labeled RNA from total, CD45^-^ or CD45^+^ cells isolated from rectal tumor (T1) and paired normal tissue (N1) (1 of 6 pairs shown). **g**, Heatmap of Sialylated and Sialofucosylated glycan abundance Z-scores (total glycan-normalized) from small RNA and membrane proteins in rectal tumor *vs.* adjacent normal tissues (n=5). **h**, Relative abundance of 5 glycan subtypes in tumor *vs.* adjacent normal tissue-derived glycoRNAs and glycoproteins. **i**, Differential *vs.* non-differential structurally defined glycans in small RNA and membrane proteins (rectal tumor *vs*. adjacent normal tissues).

To further investigate whether observed sialoglycoRNA overexpression in tumors involves specific glycoform alterations, we performed N-glycan profiling of small RNA from five pairs of rectal tumor and adjacent normal tissues using mass spectrometry, with membrane proteins from the same samples for comparison (Extended Data Fig. 3a,b). For glycoRNA, tumor tissues showed increased sialylation compared to adjacent normal tissues, consistent with the results of StCEL-based sialoglycoRNA detection (Fig. 3e,g,h and Extended Data Fig. 3c). Notably, five structurally defined sialylated glycans were significantly elevated in tumor tissues, while increases in several structurally defined fucosylated glycans occurred concurrently with marked reductions in structurally defined high-mannose glycans (Fig. 3g,h and Extended Data Fig. 3c,d). In contrast, glycoproteins barely exhibited significant changes in glycan subtypes or structurally defined glycans across matched tissue pairs (Fig. 3g-i and Extended Data Fig. 3d-f). These findings demonstrate tumor-specific glycosylation dysregulation as a hallmark of glycoRNAs, highlighting their potential as distinct biomarkers over glycoproteins.

### StCEL reveals clinical significance of sialoglycoRNA dysregulation in tissues and serum cohorts

The high expression of sialoglycoRNA in cancer prompted us to evaluate its associations with clinicopathological features. To overcome gel electrophoresis’ semi-quantitative and low-throughput limitations, we substituted dibenzocyclooctyne-biotin (DBCO-biotin) with DBCO-Cy5 in StCEL’s strain-promoted azido-alkyne cycloaddition (SPAAC). This enables direct fluorescent quantification via a microplate reader, providing quantitative data while reducing runtime (Fig. 4a). First, we validated the feasibility of incorporating DBCO-Cy5 into the StCEL workflow, which showed that DBCO-Cy5 provided the specific fluorescent signals for both gel electrophoresis and microplate reader (Extended Data Fig. 4a). Furthermore, the validation in StCEL with the analysis of serially diluted RNAs demonstrated a linear correlation (R²=0.9995) between fluorescence intensity and RNA quantity, demonstrating the accurate and robust quantification of sialoglycoRNAs by DBCO-Cy5-based StCEL (Fig. 4b and Extended Data Fig. 4b). Additionally, comparable to gel electrophoresis, the microplate reader detected distinct signals at the lowest RNA input, indicating the StCEL combined with microplate reader achieves comparable sensitivity to gel electrophoresis (Fig. 4b and Extended Data Fig. 4b).

**Fig. 4.**
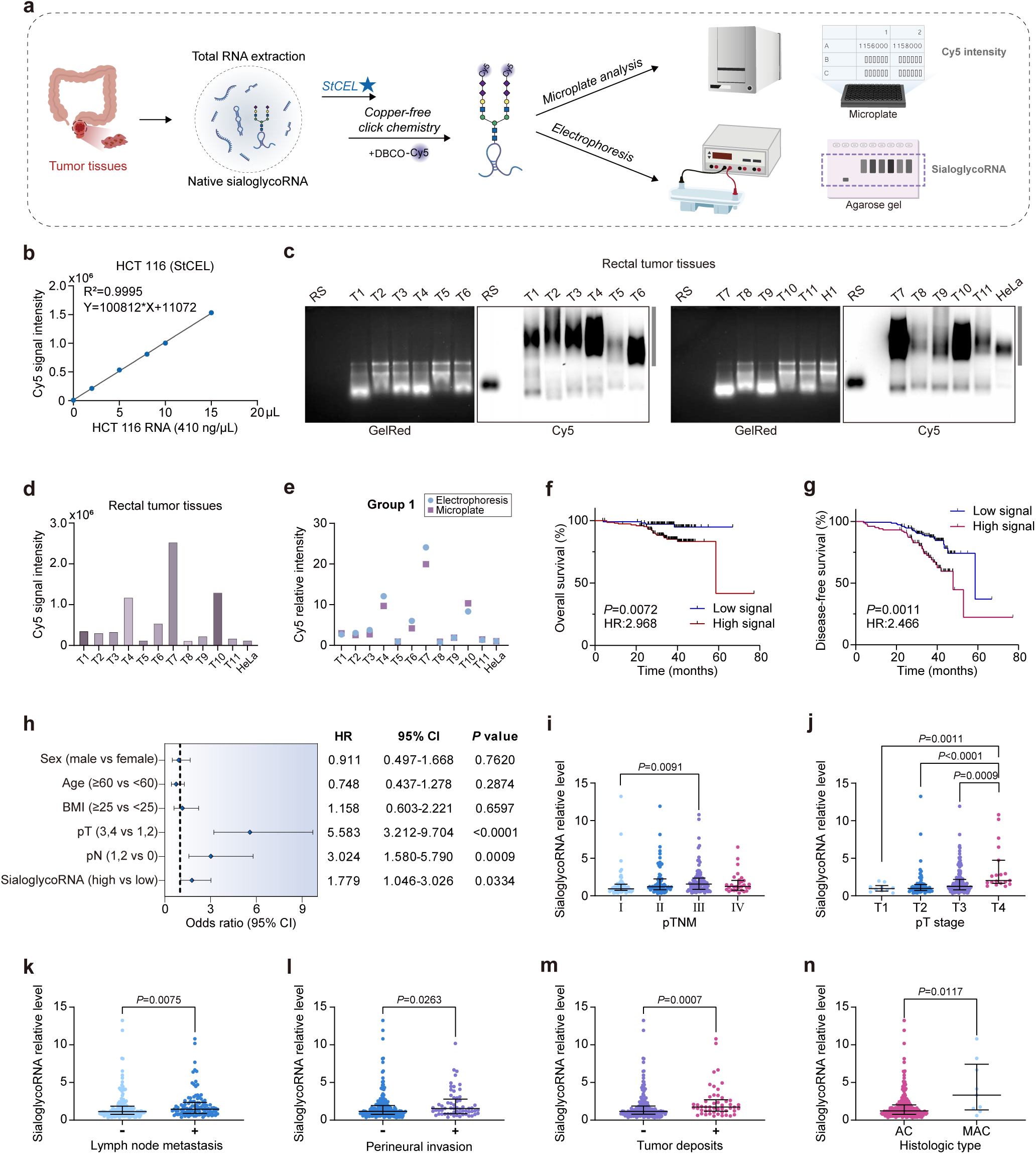
StCEL reveals the clinical significance of sialoglycoRNA dysregulation in tissues. a, Workflow for large-scale detection in rectal tumor tissues: the extracted total RNA was subjected to StCEL and SPAAC with DBCO-Cy5, followed by gel electrophoresis or microplate reading (Cy5 channel). **b**, Linear correlation between the signal values detected by the microplate reader and the amount of StCEL-labeled RNA (Cy5). **c**, RNA fluorescent imaging of sialoglycoRNA labeled by StCEL from rectal tumor tissues (representative 11 of 256 samples shown) using a multifunctional imager after gel electrophoresis. The result of the remaining tissues is shown in the **Supplementary Figure. 1e.** “RS” referred to the synthetic RNA standard (RNA-Cy5) as used in **Extended Data Fig. 4c**. “HeLa” referred to the total RNA extracted from HeLa cells, which was used to correct the labeling efficiency of different groups. **d**, Microplate reader-based quantification of the same samples as in **c. e**, Concordance validation between gel electrophoresis (**c**) and microplate reader (**d**) in representative Group 1. **f,g,** Kaplan–Meier survival curves showing overall survival (OS, **f**) and disease-free survival (DFS, **g**) of patients stratified by sialoglycoRNA expression levels. **h**, Multivariate Cox regression survival analysis (backward LR method) showed DFS in patients stratified by sialoglycoRNA expression levels. **i**, SialoglycoRNA levels across pTNM stages (Ⅰ-Ⅳ). Statistical comparisons by Kruskal-Wallis test, *P*=0.0176. **j**, SialoglycoRNA levels across pT substages (T1-T4). Statistical comparisons by Kruskal-Wallis test, *P*<0.0001. **k**, SialoglycoRNA levels stratified by lymph node metastasis status. Statistical comparisons by the Mann-Whitney test. **l**, SialoglycoRNA levels stratified by perineural invasion status. Statistical comparisons by the Mann-Whitney test. **m**, SialoglycoRNA levels stratified by tumor deposit status. Statistical comparisons by the Mann-Whitney test. **n**, SialoglycoRNA levels in adenocarcinoma (AC) vs. mucinous adenocarcinoma (MAC). Statistical comparisons by the Mann-Whitney test.

To examine the sialoglycoRNA levels in a large cohort, a synthetic RNA-Cy5 was used as a standard to correct the detection biases among different microplates (Extended Data Fig. 4c). Here, HeLa total RNA was included as an internal control in each batch to normalize technical variations across experimental batches. Strong concordance between microplate reader and gel electrophoresis imaging validated the viability of DBCO-Cy5-based detection via microplate reader (Fig. 4c-e, Extended Data Fig. 4d, and Supplementary Fig. 1e). In a cohort of 265 patients with rectal cancer, elevated sialoglycoRNA was significantly correlated with worse overall survival (OS; hazard ratio [HR] =2.968 [95% confidence interval {CI} 1.342–6.564], *P*=0.0072; Fig. 4f) and disease-free survival (DFS; HR=2.466 [95% CI 1.436–4.235], *P*=0.0011; Fig. 4g). Multivariate analysis identified sialoglycoRNA was an independent prognostic factor for DFS (HR=1.779 [95% CI 1.046–3.026], *P*=0.0334; Fig. 4h). Furthermore, the sialoglycoRNA levels was positively correlated with advanced pathological TNM (pTNM; III vs I: *P*=0.0091), higher pathological T (pT) stage (T4 vs T1: *P*=0.0011; T4 vs T2: *P*<0.0001; T4 vs T3: *P*=0.0009), lymph node metastasis (*P*=0.0075), perineural invasion (*P*=0.0263), tumor deposits (*P*=0.0007), as well as mucinous adenocarcinoma (*P*=0.0117) (Fig. 4i-n and Extended Data Fig. 4e-h). These results suggest that elevated sialoglycoRNA levels are associated with malignant potential and poor prognosis, serving as a novel biomarker for cancer progression.

Furthermore, we examined whether the StCEL could be applied for serum-derived sialoglycoRNA detection. As expected, sialoglycoRNA was also detected in human serum by StCEL, showing sensitivity to neuraminidase and RNase treatment (Extended Data Fig. 5a-c). Most importantly, sialoglycoRNA levels in the serum of patients with rectal cancer were significantly higher than those in healthy donors (Extended Data Fig. 5d), indicating a potential application in liquid biopsy.

### StCEL-based RNA-seq reveals tumor-associated dysregulation of sialoglycoRNAs

Besides detecting the expression level of sialoglycoRNA, StCEL-mediated biotin labeling also enables enrichment of sialoglycoRNA, which can be extended for high-throughput sequencing. We integrated StCEL with PANDORA-seq, a method optimized for sequencing modified small RNAs, to achieve more extensive and accurate identification^31^ (Fig. 5a). StCEL-based sialoglycoRNA-seq of HCT 116 showed that sialoglycoRNAs consisted of small RNAs less than 50 nucleotide (nt) in length, with an inverse correlation between size and abundance, mainly including rRNA-derived small RNA (rsRNA) and tRNA-derived small RNA (tsRNA; Fig. 5b and Extended Data Fig. 6a). In line with previous studies^12^, plotting StCEL-enriched RNAs against input small RNAs revealed two distinct sequence clusters (dashed red/green boxes; Fig. 5c), confirming that RNAs are specifically chosen for glycosylation. Similar results were observed with Ac_4_ManNAz-based sialoglycoRNA-seq (Extended Data Fig. 6b,c). Additionally, correlation analysis showed high consistency between StCEL-and Ac_4_ManNAz-enriched sialoglycoRNA sequencing abundances (Fig. 5d and Extended Data Fig. 6d,e), demonstrating that StCEL-based sialoglycoRNA-seq is an effective and specific method for analyzing sialoglycoRNA expression profiles.

**Fig. 5.**
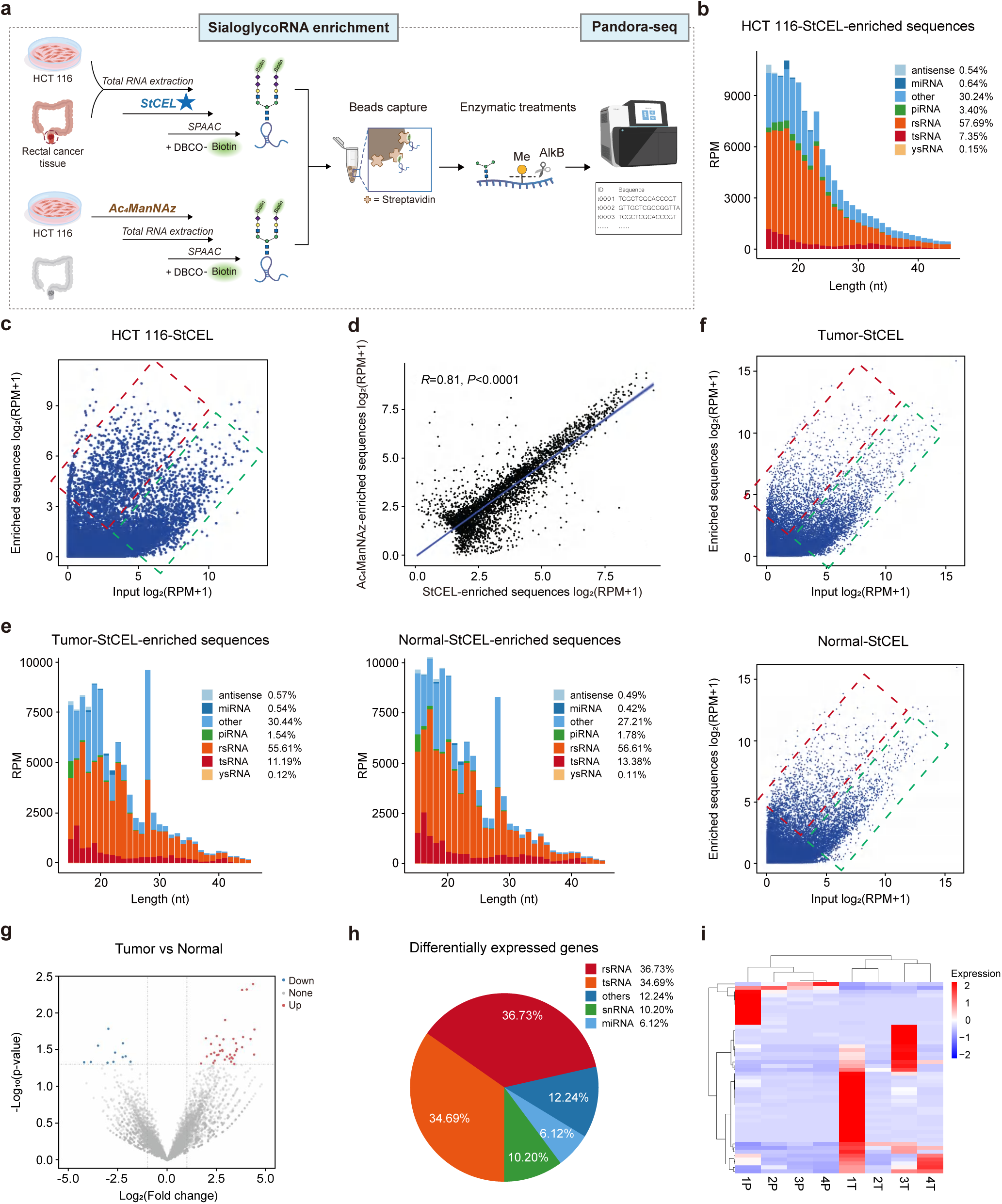
Elucidating sequence characteristics of sialoglycoRNA in cultured cells, tumor tissues, and adjacent normal tissues using StCEL. a, Workflow of sialoglycoRNA-seq using Ac_4_ManNAz-or StCEL-labeling to enrich sialoglycoRNA followed by PANDORA-seq. **b**, Length distribution of StCEL-enriched sialoglycoRNA in HCT 116 cells. **c**, Scatter plot comparing RNA sequence abundance in StCEL-purified sialoglycoRNA versus input small RNA. Red dashed boxes denote sequences enriched in purified sialoglycoRNA samples with positive abundance correlation, while green boxes indicate depleted sequences. **d**, Transcript abundance correlation: StCEL-*vs.* Ac_4_ManNAz-enriched sialoglycoRNA-seq. **e**, Length distribution of StCEL-enriched sialoglycoRNAs in 5 paired rectal tumor (left) *vs.* adjacent normal tissues (right). **f,** Scatter plots comparing RNA sequence abundance in StCEL-purified sialoglycoRNA versus input small RNA in tumor (top) or normal tissues (bottom). Red dashed boxes denote sequences enriched in purified sialoglycoRNA samples with positive abundance correlation, while green boxes indicate depleted sequences. **g**, Volcano plot of differentially enriched sialoglycoRNAs between rectal tumor and adjacent normal tissues. Red dots: significantly enriched in rectal tumor samples (log_2_ fold change [FC] > 1, adjusted p-value < 0.05). Blue dots: significantly enriched in adjacent normal samples (log_2_ FC <-1). **h**, Composition of differentially expressed genes identified in **g**. **i,** Heatmap of the differentially expressed sequences in **h**.

Given the tumor-associated upregulation of sialoglycoRNA and dysregulation of its glycan moieties, we aimed to investigate potential alterations in the RNA components between tumor and adjacent normal tissues. Therefore, we performed StCEL-based sialoglycoRNA-seq in four paired rectal tumor and adjacent normal tissues to decipher inter-tissue sequence heterogeneity. Consistent with the cellular findings, the sialoglycoRNA profiles in both rectal tumor and adjacent normal tissues were dominated by rsRNA and tsRNA (Fig. 5e and Extended Data Fig. 7a). Interestingly, unlike sialoglycoRNAs in cell lines, tissue-derived sialoglycoRNAs exhibited a pronounced enrichment at 28 nt, primarily comprising rsRNA (Fig. 5e and Extended Data Fig. 7a). Consistent with the observations in cell lines, comparative analysis of the RNA abundance in StCEL-enriched fractions versus input small RNAs revealed two distinct clusters in both rectal tumor and adjacent normal tissues, confirming selective glycosylation of RNA. (Fig. 5f and Extended Data Fig. 7b). Importantly, comparative analysis of the sialoglycoRNA abundance between rectal tumor and adjacent normal tissues revealed that some sequences were significantly dysregulated (Fig. 5g-i), most of which were identified as rsRNA and tsRNA (Fig. 5h). Additionally, the levels of sialoglycoRNAs were positively correlated with their abundance in input small RNA (Fig. 5f), suggesting global acceleration of RNA glycosylation.

## Discussion

In this study, we developed StCEL, a novel method designed to label sialoglycoRNA for detection and high-throughput sequencing. StCEL utilized monosialyltransferase CstⅡ to transfer azido-modified sialic acid to the natural sialic acid on sialoglycoRNA, enabling subsequent click chemistry-based labeling and subsequent abundance analysis or enrichment for RNA-seq. Considering that StCEL labels sialoglycoRNA after RNA extraction, it can be theoretically applied to almost all sample types. Using StCEL, sialoglycoRNA levels were noted to be significantly elevated in multiple cancer types, including rectal, lung, and esophageal cancer. Moreover, a high sialoglycoRNA levels were significantly correlated with advanced pTNM stages, lymph node metastasis, and poor survival in patients with rectal cancer. Interestingly, sialoglycoRNA signal was also detected in serum, and the serum sialoglycoRNA levels of patients with rectal cancer were higher than that of healthy donors, indicating its potential as a biomarker for cancer screening. Most significantly, StCEL-based sialoglycoRNA-seq profiling of clinical tissues revealed tumor-associated dysregulation in specific sialoglycoRNA subsets, suggesting their potential involvement in malignant progression.

StCEL offers several advantages. First, by employing post-extraction labeling, StCEL bypasses key limitations of metabolic labeling (e.g., temporal constraints and cellular toxicity) while avoiding interference from complex stress conditions, such as oxidative stress or ionizing radiation (Extended Data Fig. 8a-d). Moreover, this approach theoretically permits sialoglycoRNA detection in nearly all sample types, as previously discussed. Second, StCEL exhibits over 50-fold sensitivity compared to Ac_4_ManNAz-based labeling. Third, the sialyltransferase used in StCEL has no impact on the RNA backbone, avoiding the introduction of ribonucleic acid alterations observed in chemistry-based labeling methods like rPAL^11^. Additionally, the high specificity of the enzyme-mediated biochemical process confirmed the high specificity of StCEL, highlighting its application in high-throughput sequencing. Finally, StCEL-mediated fluorescence labeling could be detected using a microplate reader, facilitating large-scale analyses. This methodology significantly advances clinical application of sialoglycoRNA.

Interestingly, gel electrophoresis revealed distinct sialoglycoRNA band patterns in the tissue and serum samples, inconsistent with the single band of cell lines (Fig. 2j and Fig. 3b-d). Specifically, tumor tissue displayed prominent bands between the loading well and 28S rRNA. In contrast, adjacent normal tissue exhibited a weak band in this range, except for a prominent band corresponding to 5S rRNA. (Fig. 3b-d, Extended Data Fig. 2d-f and Supplementary Fig. 1a-c). Similarly, serum sialoglycoRNA displayed a dominant band near the 5S rRNA (Extended Data Fig. 5a-d). This pattern is consistent with previous studies, demonstrating that sialoglycoRNA of immune cells, such as THP-1, presented two electrophoretic bands, including a band near the 5S rRNA^32^. Given the abundance of immune cells in tissues and blood^33, 34^, the lower electrophoretic band observed in both sample types may represent immune cell-derived components. Notably, the bands observed in the tumor tissues exhibited molecular weight variations between the loading well and the 28S rRNA. However, sialoglycoRNAs are small RNAs, and their RNA moieties confer limited steric hindrance in gel electrophoresis, whereas the N-glycan profile exhibited higher structural variability (Extended Data Fig. 3b,g). This suggests that differences in electrophoretic mobility arise from glycan structural heterogeneity caused by glycan modifications and sialic acid content on sialoglycoRNA.

Using StCEL, we identified significant upregulation of sialoglycoRNA across multiple cancer types. In contrast, Ma et al. reported lower levels of glycosylated U1, U35a, and Y5 RNA in breast cancer cell lines than in noncarcinomatous MCF-10A cells^14^. This divergence may arise from cell-type-specific variations in Ac_4_ManNAz metabolism or fundamental constraints in recapitulating tumor malignancy spectra with in vitro cell line models. Consistent with our results, drFRET-based detection revealed elevated glycosylated U1, U35a, U3, U8, and Y5 RNAs in serum sEVs from breast cancer patients than in healthy donors^16^, suggesting their diagnostic potential as clinical biomarkers. Notably, beyond RNA expression levels, glycoRNAs also exhibit tumor-associated glycan alterations, including increased sialylation and fucosylation glycan subtypes. Additionally, StCEL-based RNA-seq revealed specific dysregulated sialoglycoRNAs in tumor tissues. Collectively, these findings suggest that sialoglycoRNA serve as a promising biomarker in cancer biology.

Although StCEL provides greater sensitivity and specificity for sialoglycoRNA detection, its limitations require further resolution. First, StCEL only detects sialylated glycoRNA, potentially overlooking glycoRNA without sialic acids. Second, despite applying PANDORA-seq, a fraction of glycoRNA-derived reads failed to map to the reference genome, suggesting that glycan modifications may hinder reverse transcriptase processivity during glycoRNA sequencing. Treatments to remove modifications before sequencing may improve performance. In summary, the StCEL method establishes a multifunctional platform that enables sensitive and specific detection, coupled with sequencing of sialoglycoRNA, highlighting its significant potential for clinical applications.

## Methods

### Cell culture

Human cells were cultured in an incubator with 5% CO_2_ at 37°C. HeLa (American Type Culture Collection [ATCC]), RKO (ATCC), MDA-MB-231 (ATCC), and HEK293T (ATCC) cells were cultured in DMEM medium (Gibco, California, USA; Cat. No. C11995500BT) supplemented with 10% fetal bovine serum (FBS; Corning, New York, USA; Cat. No. 35-179-CV) and 1% penicillin/streptomycin (Gibco; Cat. No. 15140-122). HCT 116 (ATCC) and NCI-H460 (ATCC) cells were cultured in RPMI-1640 medium (Gibco; Cat. No. C11875500BT) supplemented with 10% FBS and 1% P/S.

Insect cells Sf9 (ATCC) and Hi-Five (Thermo Fisher Scientific, Waltham, Massachusetts, USA; Cat. No. B85502) were cultured in Grace’s Insect Medium supplemented with 10% FBS and 1% P/S in an incubator at 27℃ with room air.

### The expression and purification of CstⅡ

The monosialyltransferase CstⅡ was produced by *E. coli* expression system. Briefly, the coding sequence of CstⅡΔ32I53S with a 6×His tag was cloned into the pTriEx vector (pTriEx-CstⅡΔ32I53S). Escherichia coli strain BL21 Star™ (DE3) was transformed with the pTriEx-CstⅡΔ32I53S plasmid and cultured in LB medium containing the appropriate antibiotic at 37°C with shaking at 220 rpm. When the optical density at 600 nm (OD600) reached approximately 0.6, IPTG was added to a final concentration of 0.1 mM, and the culture was incubated further at 20°C for 32 hours. For purification, bacteria were pelleted by centrifugation, resuspended in lysis buffer (100 mM Tris-HCl, pH 8.0, 0.1% Triton X-100), and subjected to sonication. Following centrifugation, the supernatant was loaded onto a gravity-flow Ni-NTA column. The column was washed with Wash Buffer (20 mM Tris-HCl, 500 mM NaCl, 50 mM imidazole, pH 8.0), and the His-tagged protein was eluted with Elution Buffer (20 mM Tris-HCl, 500 mM NaCl, pH 8.0, containing imidazole at final concentrations of 100 mM and 500 mM). The eluted protein was then loaded onto a Chromdex 200 column with running buffer (50 mM Tris-HCl, 500 mM NaCl, 10% glycerol, pH 8.0). The purified protein was sterilized using a 0.22 μm filter before being aliquoted for storage. The purity of the recombinant protein was evaluated by SDS-PAGE with Coomassie Brilliant Blue staining, demonstrating > 95% homogeneity.

### In vitro transcription

In vitro transcribed RNA was produced using the T7 High Yield RNA Transcription kit (Vazyme Biotech, Nanjing, China; Cat. No. DD4202-01) according to the manufacturer’s instructions. Briefly, 2 μL of Control DNA Template (0.5 μg/μL, provided in the kit) was combined with 2 μL of 10× Reaction Buffer, 2 μL of ATP Solution, 2 μL of GTP Solution, 2 μL of UTP Solution, 2 μL of CTP Solution, 2 μL of T7 RNA Polymerase Mix, and 8 μL of RNase-free H_2_O, then incubated at 37°C for 12 hours. The resulting RNA was purified with TRIzol (Thermo Fisher Scientific; Cat. No. 15596026) reagent.

### Metabolic chemical reporters and inhibitors

Stock of Ac_4_ManNAz (N-azidoacetyl-mannosamine-tetraacylated; MCE [MedChemExpress], New Jersey, USA; Cat. No. HY-W728531) was made to 500 mM in DMSO. For cell treatment, Ac_4_ManNAz was used at a final concentration of 100 μM. Stock of CMP-Neu5Az (Qiyuebio, Xi’an, China; Cat. No. Q-0069741) was made to 25 mM in 100 mM Tris HCl (pH 6.6). Stocks of glycan-biosynthesis inhibitors were all made in DMSO at the following concentrations and stored at-80℃: 10 mM NGI-1 (MCE; Cat. No. HY-117383), 10 mM kifunensine (MCE; Cat. No. HY-19332), 50 mM P-3F_AX_-Neu5Ac (MCE; Cat. No. HY-110288).

### Clinical tissues and serum, along with mouse tissues

Fresh rectal tumor tissues (for CD45⁻/CD45⁺ cell isolation) and additional tumor specimens (rectal, lung, esophageal) were sourced from the First Affiliated Hospital of Zhengzhou University under institutional ethics approval. The resected tissues were immediately placed on ice and transferred to a-80°C freezer for long-term storage. For serum fractionation, the patient’s blood was kept at room temperature for 1 hour, centrifuged at 3,000 rpm for 15 min at 4°C, and the supernatant (serum) was transferred to a new tube and stored at-80°C. The study protocol complied with the Declaration of Helsinki and institutional guidelines of Zhengzhou University and was approved by the ethics committees of the First Affiliated Hospital of Zhengzhou University (2024-KY-1497-002).

The animal experiments were approved by the Institutional Animal Care and Use Committee of Zhengzhou University (Accreditation No. ZZU-LAC20230804[10]). Experimental procedures were performed under the Guide for the Care and Use of Laboratory Animals (National Institutes of Health Publication No. 80-23, revised 1996) and according to the institutional ethical guidelines for animal experiments.

### Isolation of CD45^+^ and CD45^-^ cells from fresh human tissues

Fresh human tumor and adjacent normal tissues were mechanically dissociated and enzymatically digested to generate single-cell suspensions, which were filtered through a 70 μm cell strainer (Biosharp Life Sciences, Hefei, CN; Cat. No. BS-70-XBS) and washed in ice-cold sorting buffer (PBS, pH 7.2, containing 0.5% BSA and 2 mM EDTA). CD45^+^ and CD45^-^ cells were then isolated from these suspensions using the Human CD45^+^ Cell Isolation kit (RWD Life Science, Shenzhen, CN; Cat. No. K1205-10) coupled with LarSep Columns (RWD Life Science; Cat. No. HCSC-25). Briefly, cells were adjusted to ≤1×10⁸/mL in buffer, incubated with 10 μL/mL biotinylated anti-human CD45 antibody (provided in the kit) for 10 min at 4°C, washed, and subsequently incubated with 10 μL/mL streptavidin-conjugated magnetic beads (provided in the kit) for 15 min at 4°C. After washing, labeled cells were resuspended in buffer (500 μL-1 mL per ≤1.25×10⁸ cells), loaded onto a pre-washed (2 mL buffer) magnetic column, and unlabeled (CD45^−^) cells were collected in the flow-through following 2-3 column washes. Bound CD45^+^ cells were eluted with 1-2 mL buffer after column removal from the magnet. All procedures were performed under sterile, cold (4°C) conditions to minimize non-specific binding.

### RNA extraction and purification

Homogenized tissues or cell lines were lysed with TRIzol, mixed with 0.2 volumes of chloroform, vortexed thoroughly, and centrifuged at 12,000 ×g for 15 min at 4℃. The upper layer was transferred to a fresh tube, mixed with 1 volume of isopropanol, incubated at room temperature, and centrifuged at 12,000 ×g for 30 min at 4℃. The RNA pellet was washed with pre-chilled 70% ethanol and then dissolved in nuclease-free water. For serum samples, 750 μL TRIzol was added to 250 μL serum, vortexed to mix, and the RNA extraction is described above. To remove the residual protein, RNA was mixed with Proteinase K (Pro K, Thermo Fisher Scientific; Cat. No. AM2546) to a final concentration of 1 μg per 1μg RNA and incubated for 45 minutes at 37℃, followed by RNA purification with TRIzol as described above.

The RNA purification after labeling or treatment was performed using Zymo RNA Clean & Concentrator™ kit (Zymo Research, Irvine, California, USA; Cat. No. R1018). Briefly, RNA mixture was added with 2 volumes of RNA binding buffer (example: mix 100 µL buffer and 50 µl sample), vortexed, and mixed with an equal volume of ethanol (example: add 150 µL ethanol). Then, the mixture was transferred to Zymo columns and centrifuged at 12,000 ×g for 30 seconds, washed sequentially with 400 μL RNA Prep Buffer and 400 μL RNA Wash Buffer twice. Finally, 25 μL (Zymo-Spin™ IICR Columns, provided in the kit) or 50 μL (Zymo-Spin™ IIIGC Columns; Cat. No. C1006-250-G) nuclease-free water was added to the column and centrifuged at 12,000 × g for 30 seconds to collect eluted RNA.

### Small and large RNA fractionation

Small and large RNAs were separated using the Zymo RNA Clean & Concentrator™ kit. Briefly, the extracted total RNA was mixed with an equal 2 volumes of adjusted RNA Binding Buffer (Mix an equal volume of RNA Binding Buffer and 100% ethanol) and loaded onto a silica column. After centrifuging at 12,000 ×g for 30 seconds, the flow-through containing small RNA (<200 nt) was collected. The collected RNA was then mixed 1:1 with ethanol and purified as described above. And the large RNA binding to the column was purified as described above.

### StCEL

The reaction mixture was prepared by sequentially adding the following components to the extracted total RNA: CstⅡ, CMP-Neu5Az, Mg²⁺, and Tris-HCl. The mixture was incubated at 37°C for 3 hours. StCEL-labeled RNA was purified using the Zymo RNA Clean & Concentrator™ kit as described. Next, to perform copper-free click conjugation with sialoglycoRNA, DBCO-PEG4-Biotin (Merck, Darmstadt, Germany; Cat. No. 760749) or DBCO-Sulfo-Cy5 (Merck; Cat. No. 777374) was employed as the cyclooctyne component for the SPAAC reaction. Briefly, the SPAAC reaction mixture consisted of 10 μL “dye-free” Gel Loading Buffer II (df-GLBII, 95% formamide, 18mM EDTA, and 0.025% SDS), 9 μL RNA, and 1 μL DBCO reagent (10 mM in stock). After incubation at 55℃ for 10 min, reactions were quenched by adding 80 μL nuclease-free water, followed by purification using the Zymo column as described above and analyzed by gel electrophoresis or a microplate reader.

### Enzymatic treatment of RNA samples

Multiple enzymes, such as endonucleases, exonucleases, and neuraminidase, were utilized for substrate digestion. Specifically, nucleases targeted RNA/DNA, while neuraminidase acted on sialic acid residues. All digestions were performed on 20 μg of total RNA in a 20 μL reaction volume at 37℃ for 60 minutes. To digest RNA, the following enzymes were used: 2 μL RNase cocktail (containing 0.5U/μL RNase A and 20U/μL RNase T1; Thermo Fisher Scientific; Cat. No. AM2286) in 1×TURBO DNase buffer (provided in the kit of Cat. No. AM2238) or 2 μL RNase I (Thermo Fisher Scientific; Cat. No. EN0602) in 1 ×RNase I buffer (20 mM Tris-HCl [pH 8.0], 100 mM NaCl, 0.1 mM EDTA, 0.01% Triton X-100). To digest DNA, 2 μL TURBO DNase (2 U/μL, Thermo Fisher Scientific; Cat. No. AM2238) was used in 1 × TURBO DNase buffer. To digest sialic acid residues, 2 μL α2-3,6,8,9 Neuraminidase A (50 U/μL; NEB; Cat. No. P0722) was used in 1 × GlycoBuffer 1 (provided in the kit).

### RNA gel electrophoresis, blotting, and imaging

RNA (purified, enriched, or enzymatically digested) was purified using the Zymo column and finally eluted with 25 μl of df-GLBII. After incubating at 55℃ for 10 minutes and chilling on ice for 3 minutes, RNA was then loaded onto a 1% agarose-formaldehyde denaturing gel stained with GelRed (1:20000; Biosharp Life Sciences; Cat. No. BS354B) and electrophoresed at 115 V for 30 minutes. Total RNA was then visualized in the gel using a UV gel imager (BLT, Guangzhou, CN; GelView 5000Pro Ⅱ). After being transferred to a 0.45 mm nylon membrane (Cytiva Life Sciences, Washington, D.C., USA; Cat. No. RPN303B), RNA was crosslinked to the nylon membrane using UV-C light (0.18 J/cm^2^). Membrane was then blocked with 1×Blocking Buffer (Beyotime, Shanghai, CN; Cat. No. D3308B) for 15 minutes, incubated with Streptavidin-HRP (1:2000; Beyotime; Cat. No. A0303) in 1×Blocking Buffer for 15 minutes, washed with 1×Washing Buffer (Beyotime; Cat. No. D3308W) for three times at room temperature, and balanced in 1×TBST Buffer. Then, the membrane was developed using chemiluminescent substrate (Merck; Cat. No. WBKLS0500) in the Amersham ImageQuant 800 (Cytiva). Images were quantified with the ImageJ software.

### Western blotting

Whole-cell lysates were prepared using RIPA Buffer (NCM, Suzhou, CN; Cat. No. WB3100) supplemented with EDTA-free Protease Inhibitor Cocktail (Merck; Cat. No. 539134). Proteins were separated by FuturePAGE™ precast gels (ACE, Suzhou, CN; Cat. No. ET10420Gel) and transferred to PVDF membranes (Merck; Cat. No. IPVH00010) using a wet transfer system. Membranes were blocked with 1×TBST containing 5% milk (Biosharp Life Sciences; Cat. No. BS102-500g) for 1 hour at room temperature and incubated with primary antibodies overnight at 4℃. The primary antibodies and concentrations used for Western blotting were as follows: anti-Na^+^/K^+^-ATPase (1:2000, HUABIO, Hangzhou, CN; Cat. No. ER64104), anti-Lamin A/C (1:2000, HUABIO; Cat. No. ET7110-12), and anti-GAPDH (1:10000, Proteintech, Wuhan, CN; Cat. No. 60004-1-Ig). Following incubation with HRP-conjugated secondary antibodies (Liankebio, Hangzhou, CN; Cat. No. GAR007 and GAM0072), membranes were imaged using chemiluminescent substrate in the Amersham ImageQuant 800.

### Subcellular fractionation

Cytoplasmic and nucleoplasmic cell fractions were isolated from cells as described previously^35^. Following fractionation, RNA was extracted using TRIzol reagent. The efficacy of subcellular fractionations was verified by Western blotting analysis of compartment-specific protein markers (e.g., Lamin A/C for nucleoplasm, GAPDH and Na^+^/K^+^ ATPase for cytoplasm).

### Enrichment of SialoglycoRNA

After Ac_4_ManNAz-or StCEL-mediated biotin labeling of cellular RNA, small RNA was isolated using a Zymo RNA Clean & Concentrator™ kit. SialoglycoRNA was then enriched and purified with streptavidin-conjugated beads^36^. Specifically, 10 μL of MyOne™ C1 Streptavidin beads (Thermo Fisher Scientific; Cat. No. 65001) were blocked with 50 ng/μL glycogen (Thermo Fisher Scientific; Cat. No. AM9516) in Biotin Wash Buffer (10 mM Tris-HCl pH 7.5, 1 mM EDTA, 100 mM NaCl, 0.05% Tween-20) for 1 hour at room temperature before incubation with the biotinylated sialoglycoRNA. Next, 10 μg of labeled small RNA was diluted in 750 μL of Biotin Wash Buffer and mixed with the blocked MyOne C1 beads for 2 hours at 4°C. After washing twice with 1 mL of ChIRP Wash Buffer (2×SSC, 0.5% SDS), twice with 1 mL of Biotin Wash Buffer, and twice with NT2 Buffer (50 mM Tris-HCl, pH 7.5, 150 mM NaCl, 1 mM MgCl_2_, 0.005% NP-40) for 3 minutes each, the biotinylated sialoglycoRNA was released from the beads by adding TRIzol reagent and heating at 70°C for 10 minutes, followed by RNA extraction as described above.

### PANDORA-seq of sialoglycoRNA from cells and tissues

PANDORA-seq was performed according to the protocol^31^ by Guangzhou Epibiotek Co., Ltd. (Guangzhou, CN). Briefly, the library was prepared by QIAseq® miRNA Library kit (Qiagen, Hilden, Germany; Cat. No. 331505). The software SPORTS1.1 was employed for small RNA sequence annotation. Detailed information is also described as follows. Small RNA was isolated as described above. To overcome biases introduced by RNA modifications, extracted RNA underwent sequential enzymatic treatments. First, RNA was demethylated via incubation with 4 μg/mL AlkB (Guangzhou Epibiotek Co., Ltd) at 37°C for 30 minutes in an optimized reaction buffer (50 mM HEPES [pH 8.0], 75 μM ferrous ammonium sulfate, 1 mM α-ketoglutaric acid, 2 mM sodium ascorbate, 2,000 U RNase inhibitor, and 50 mg/mL BSA). Subsequently, RNA was dephosphorylated using 10U T4 Polynucleotide Kinase (T4PNK; NEB; Cat. No. M0201L) at 37°C for 20 minutes in 50 μL 1×PNK buffer (NEB; Cat. No. B0201S) supplemented with 10 mM ATP (NEB; Cat. No. P0756S). Following each enzymatic treatment, RNA was re-purified using TRIzol reagent. Library construction was then performed with the QIAseq® miRNA Library kit (Qiagen), which involved adapter ligation and PCR amplification. Final libraries were sequenced on the Illumina system.

Small RNA sequencing reads were annotated using SPORTS1.1 (updated from SPORTS1.0 with mismatch tolerance parameter-M1), followed by sequential mapping to hierarchical non-coding RNA databases in priority order: (1) miRBase v21 for miRNAs, (2) Genomic tRNA Database (GtRNAdb) for tRNAs, (3) mitotRNAdb for mitochondrial tRNAs, (4) NCBI-derived nucleotide/gene databases for rRNAs/YRNAs, (5) piRBase and piRNABank for piRNAs, and (6) Ensembl/Rfam v12.3 for other ncRNAs. For tRNA-derived small RNA (tsRNA) annotation, both pre-tRNA and mature tRNA references were utilized, with mature tRNA sequences processed through: (i) removal of predicted introns, (ii) addition of 3′-CCA termini, and (iii) 5′-G nucleotide addition specifically for histidine tRNAs; tsRNAs were subsequently classified into four subtypes: 5′ tsRNA (from pre/mature tRNA 5′ end), 3′ tsRNA (from pre-tRNA 3′ end), 3′ tsRNA-CCA (from mature tRNA 3′ terminus), and internal tsRNA (non-terminal fragments). rRNA-derived small RNA (rsRNA) mapping employed an ascending-length strategy to ensure unique parent rRNA assignment, prioritizing shorter rRNAs (e.g., 5.8S rRNA over overlapping 45S regions). Differential expression analysis of small non-coding RNAs was performed using DESeq2 v1.38.3 in R v4.2.2 with significance thresholds of |log₂FC| > 1 and adjusted p-value < 0.05.

### LC-MS/MS analysis of N-glycans from tissue glycoRNA and membrane proteins

After protein and RNA extraction, small RNA and cell membrane protein samples were processed in parallel for N-glycan analysis following previously established protocols^37, 38^. Briefly, 200 μL of 100 mM HEPES buffer was added to RNA or proteins, and heated at 100℃ for 2 minutes using a dry bath. After denaturation, 4 μL of PNGase F (500,000 units/mL, NEB; Cat. No. P0704) was added to the mixture, and the sample was incubated overnight at 37℃ to release N-linked glycans. N-glycans from both small RNA and protein samples were purified using Hypercarb SPE 96-well plates (Thermo Scientific; Cat. No. 60302). For protein samples, 400 μL of water was added, and the mixture was centrifuged at 10,000 g at 4℃ for 20 minutes, followed by collecting the supernatants for purification. For RNA samples, 400 μL of water was added, and the samples were directly loaded for purification. The purification procedure involved sequential washing with 0.1% (v/v) trifluoroacetic acid (TFA) in water, followed by elution with 40% acetonitrile containing 0.05% TFA and 80% acetonitrile containing 0.1% TFA. The combined eluates were dried using a CentriVap Centrifugal Concentrator (Labconco) and reconstituted in water with 0.1% (v/v) formic acid (FA) before LC-MS/MS analysis.

For glycomic analysis, 4 µL of samples were injected using a Waters nanoAcquity UPLC M-Class system (Waters) coupled to a SCIEX ZenoTOF 7600 mass spectrometer (SCIEX). Glycan separation was performed using a Hypercarb porous graphitic carbon HPLC column (1 mm × 100 mm, 3 μm; Thermo Scientific) at a flow rate of 50 μL/min. The mobile phases included buffer A (0.1% formic acid in water) and buffer B (acetonitrile with 0.1% formic acid). The chromatographic gradient was programmed as follows: 0–2 min, 2% B; 2–10 min, 0–12% B; 10–44 min, 12–72% B; 44–45 min, 72–100% B; 45–50 min, 100% B; 50–51 min, 100–0% B; and 51–60 min, 0% B.

Mass spectrometry analysis was conducted in positive SWATH mode with the Zeno trap enabled. Instrument parameters were set as follows: ion source gas1 at 30 psi, gas2 at 40 psi, and curtain gas at 35 psi. MS1 spectra were acquired over an m/z range of 400–2000 with a 0.2 s accumulation time. Glycan fragmentation was performed using 48 SWATH windows, dynamic collision-induced dissociation, and automated collision energy settings at a charge state of two. Product ions were monitored from m/z 140 to 2000 with a 15 ms accumulation time per window. Subsequent glycan identification and quantitation were performed using GlycanDIA Finder and Skyline software^38, 39^.

## Data availability

Data are available in the main Figures, Extended Data Figures, and Supplementary Figures. Any additional requests for information can be sent to the lead contact.

## Acknowledgements

We are grateful for the moral support from Dr Lei Wang, former vice president of the Sixth Affiliated Hospital of Sun Yat-sen University, who dedicated his lifetime to colorectal cancer and radiation-induced intestinal injury. Although Dr. Lei Wang has been away from us for six years, his noteworthy contributions to human health, as well as his extraordinary benevolence, dauntlessness, and selflessness, as recorded in our previous Lancet Digital Health paper, Nucleic Acids Research, and PNAS papers, are still engraved in our minds. Dear Dr. Wang, thank you very much. We also thank Hong-Qiang Qi for critical discussions, Su-Ying Ding for the serum samples, and Xiang-Nan Li for providing the Esophageal tumor and adjacent normal tissues. This study was supported by the National Science Fund for Distinguished Young Scholars (No. 82225040), the National Key R&D Program of China (No. 2022YFC2503700 and No. 2022YFA1105300), the National Natural Science Foundation of China (No. 82122057 and No.82373513), the Science and Technology R&D program of Henan province (No. 235200810091), and the “Three 100” Program of Henan Academy of Innovations in Medical Science.

## Author contributions

X.J.F., X.B.W., Y.L.W., and G.D.Z. designed the experiments. G.D.Z., Y.L.W., X.H.C., and Y.M. performed the experiments. H.L.L. and H.N. provided support for purifying protein technology.

L.Y.F. provided support for the data analysis of RNA-seq. L.Y., and J.Y.J. performed the LC-MS/MS. Z.N., M.W., X.Y.L., Y.S.C., M.B.Q., and J.S. performed the Western blot and Tissue homogenate. Z.Z., W.P.S., Y.B., Q.B.Z., and G.Q.Z. collected the rectal, lung, and esophageal tissues. G.D.Z., X.H.C., Y.M., Z.Z., and J.F.C. collected the serum. Y.L.W., Z.G.D., X.J.F., and X.B.W. wrote the manuscript. P.L., Y.X., and J.Z. revised the manuscript. P.L., X.J.F., X.B.W., and Y.X. supervised this study. All authors read and approved the final manuscript.

## Extended Data Figure legends

**Extended Data Fig. 1.**
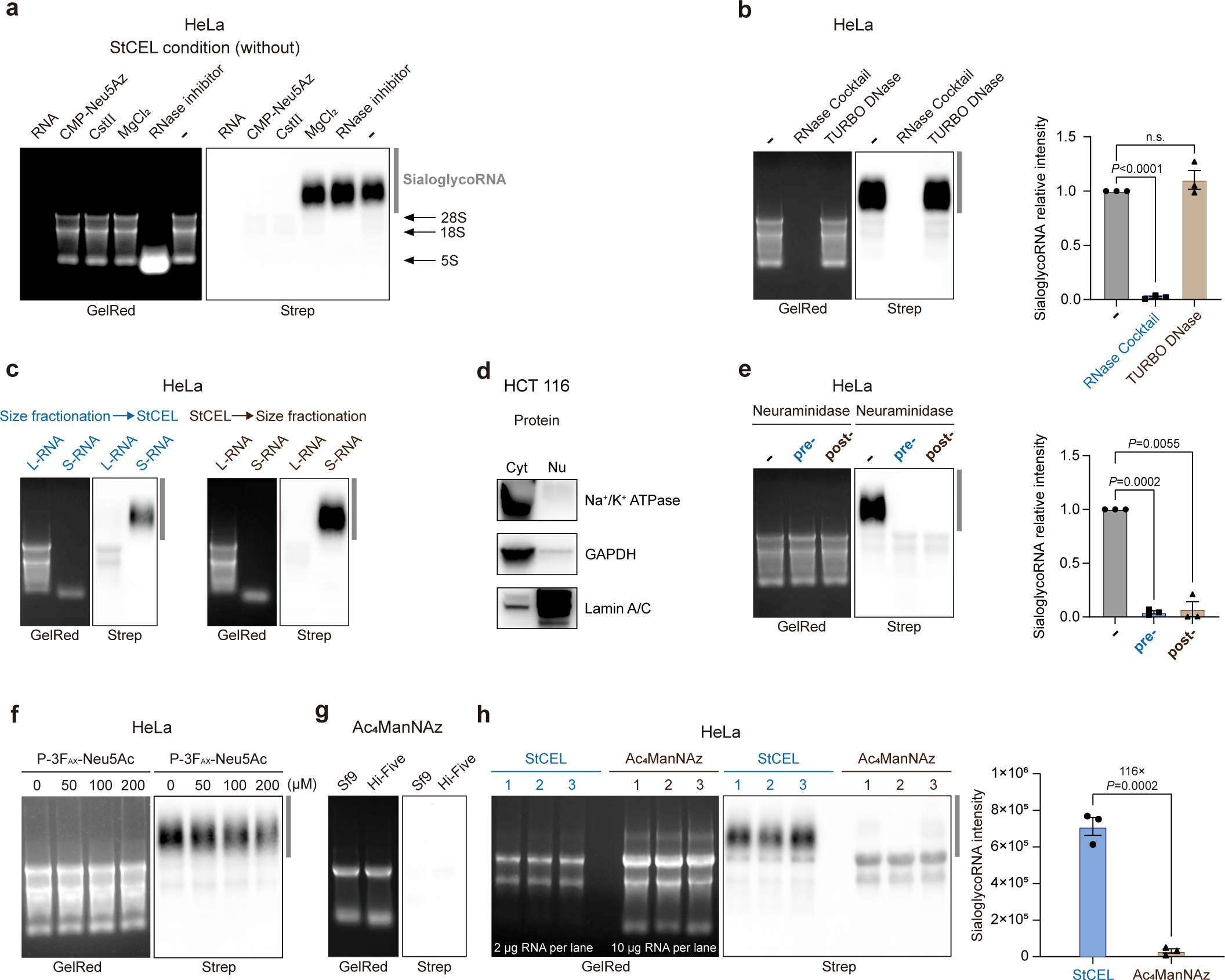
Development of StCEL and validating the sensitivity and specificity of StCEL. **a**, RNA blotting of HeLa total RNA labeled with StCEL systems in the absence of individual components (RNA, CMP-Neu5Az, CstⅡ, MgCl_2_, or RNase inhibitor). The region of sialoglycoRNA was marked by dark gray vertical bars. **b**, RNA blotting of StCEL-labeled HeLa RNA treated with RNase cocktail (A/T1) or TURBO DNase in vitro (left). Each datapoint (biological triplicate) was displayed with the mean ± SEM (right), and statistical analysis was performed with paired *t* tests. n.s. means not significant. **c**, Like the protocol in Fig. 2c, RNA blotting of HeLa large RNA (L-RNA) and small RNA (S-RNA) population of total RNA. **d,** Western Blot validated Cyt/Nu fractions from Fig. 2d using the following markers: Na^+^/K^+^ ATPase (Cyt), GAPDH (Cyt), and Lamin A/C (Nu). **e**, Like the protocol in Fig. 2e, RNA blotting of HeLa total RNA treated with neuraminidase (α2-3,6,8,9 Neuraminidase A) before or after StCEL. Each datapoint (biological triplicate) was displayed with the mean ± SEM (right), and statistical analysis was performed with paired *t* tests. **f**, HeLa cells were treated with the indicated concentrations of P-3F_AX_-Neu5Ac in complete media for 24 hours. Total RNA was then collected, labeled with StCEL, and analyzed by RNA blotting. **g**, RNA blotting of Sf9 and Hi-Five total RNA labeled with Ac_4_ManNAz. **h**, Like the protocol in Fig. 2i, RNA blotting of HeLa total RNA labeled with StCEL or Ac_4_ManNAz. The loaded amount of the StCEL-labeled group was provided in the image (left). Each datapoint (biological triplicate) is displayed with the mean ±SEM (right), and the statistical analysis was performed with an unpaired *t*-test.

**Extended Data Fig. 2.**
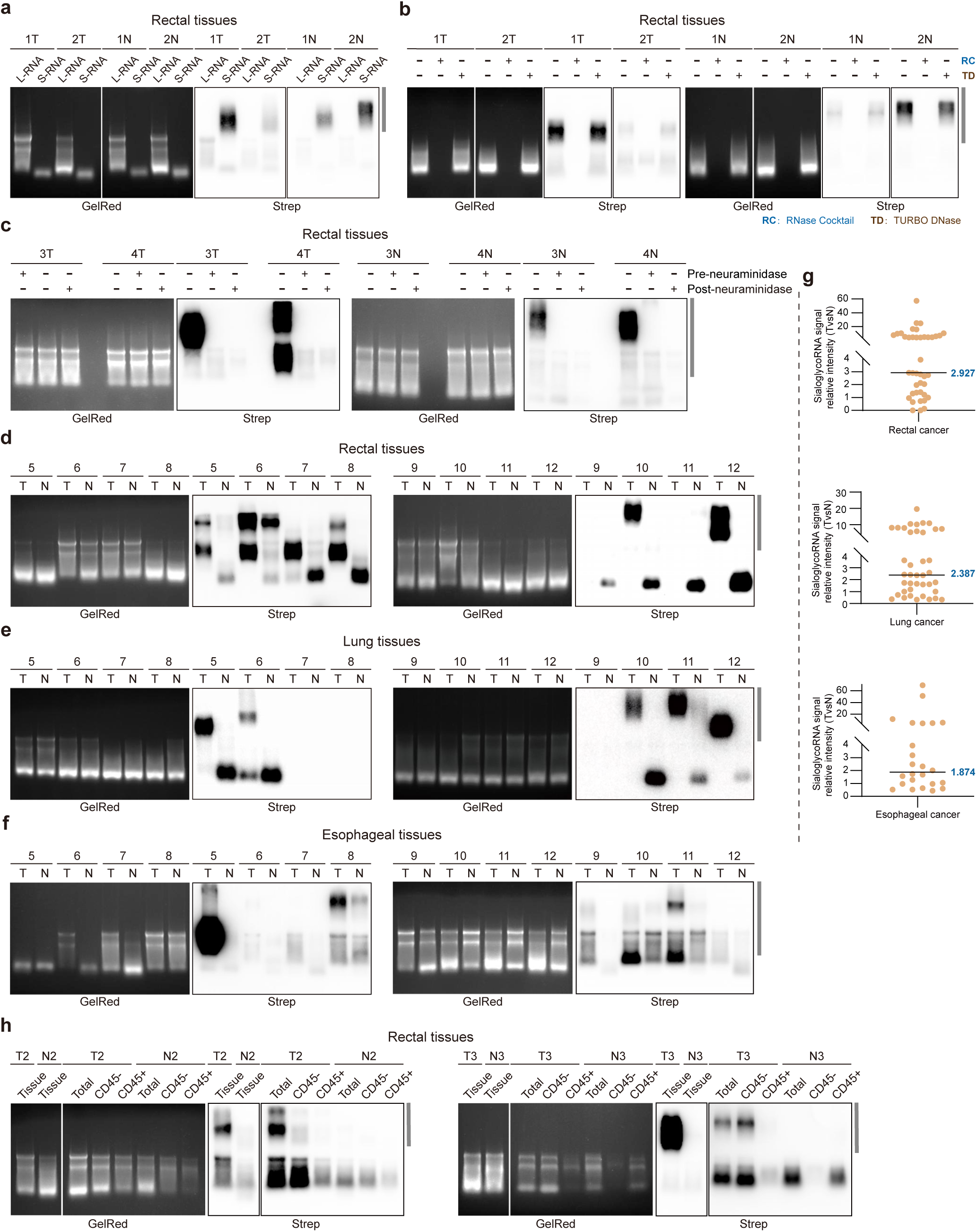
Specificity validation of StCEL for detecting tissue sialoglycoRNA and the comparison between tumor and adjacent normal tissues. **a**, RNA blotting of StCEL-labeled large RNA (L-RNA) and small RNA (S-RNA) fractions extracted from two paired rectal tumor (T) and adjacent normal (N) tissues. **b**, RNA blotting of StCEL-labeled total RNA treated with RNase cocktail (A/T1) or TURBO DNase from two paired rectal tumor (T) and adjacent normal (N) tissues. **c**, RNA blotting of total RNA treated with neuraminidase (α2-3,6,8,9 Neuraminidase A) from 2 paired rectal tumor (T) and adjacent normal (N) tissues. **d-f**, Comparative sialoglycoRNA signal intensity between tumor (T) and adjacent normal (N) tissues in **(d)** rectal cancer (representative 8 of 41 pairs shown), (**e**) lung cancer (representative 8 of 40 pairs shown), and (**f**) esophageal cancer (representative 8 of 24 pairs shown). **Supplementary Fig. 1a-c** present RNA blotting data for the remaining tissues. **g**, The expression levels of sialoglycoRNA were quantified by grayscale analysis in tumor and adjacent tissues from patients with rectal (upper panel), lung (middle panel), and esophageal (lower panel) cancer. Fold changes in tumor sialoglycoRNA expression were calculated using adjacent normal tissue expression as the baseline (normalized to 1). **h**, RNA blotting of StCEL-labeled total RNA from total tissue, CD45^-^ or CD45^+^ cells isolated from the remaining two of six pairs of rectal tumor (T) and adjacent normal (N) tissues.

**Extended Data Fig. 3.**
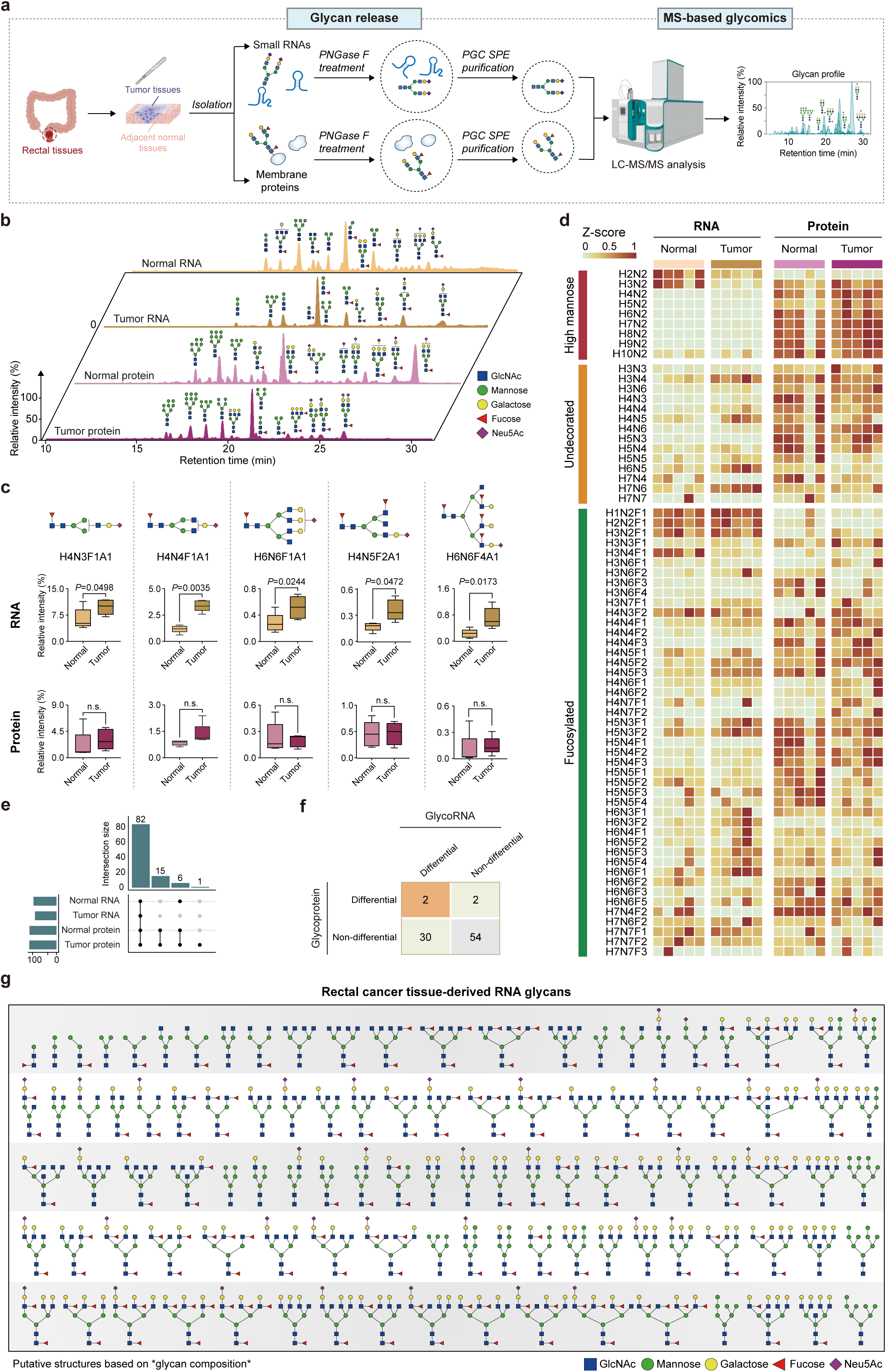
N-glycan characteristics in 5 paired rectal tumor and adjacent normal tissues. **a**, Workflow for N-glycan profiling of glycoRNAs and glycoproteins from 5 paired rectal tumor and adjacent normal tissues via LC-MS/MS: sample preparation (small RNA and membrane protein extraction), deglycosylation (PNGase F treatment), purification, and LC-MS/MS analysis. LC-MS, liquid chromatography-tandem mass spectrometry. **b**, LC-relative abundance profiles of distinct glycans from Fig. 3g and **Extended Data Fig. 3d. c**, Box plots showing relative abundance of representative N-glycan structures in glycoRNA between tumor (T) and adjacent normal (N) tissues, compared to their abundance in glycoproteins. Each datapoint is displayed with the mean ± SEM, and statistical analysis was performed by a paired *t*-test. n.s. means not significant. **d**, Heatmap of High mannose, Undecorated and Fucosylated glycans abundance Z-scores (total glycan-normalized) from small RNA and membrane proteins in rectal tumor *vs.* adjacent normal tissues (n=5). **e**, Upset plot: intersection of structurally defined glycans identified by LC-MS/MS in tumor *vs.* adjacent normal tissues (both small RNA and membrane proteins). **f**, Differential *vs.* non-differential structurally defined glycans in small RNA and membrane proteins (rectal tumor *vs*. adjacent normal tissues). **g**, Proposed glycan structures inferred from the compositional data obtained from small RNA and membrane protein of rectal tumor tissues.

**Extended Data Fig. 4.**
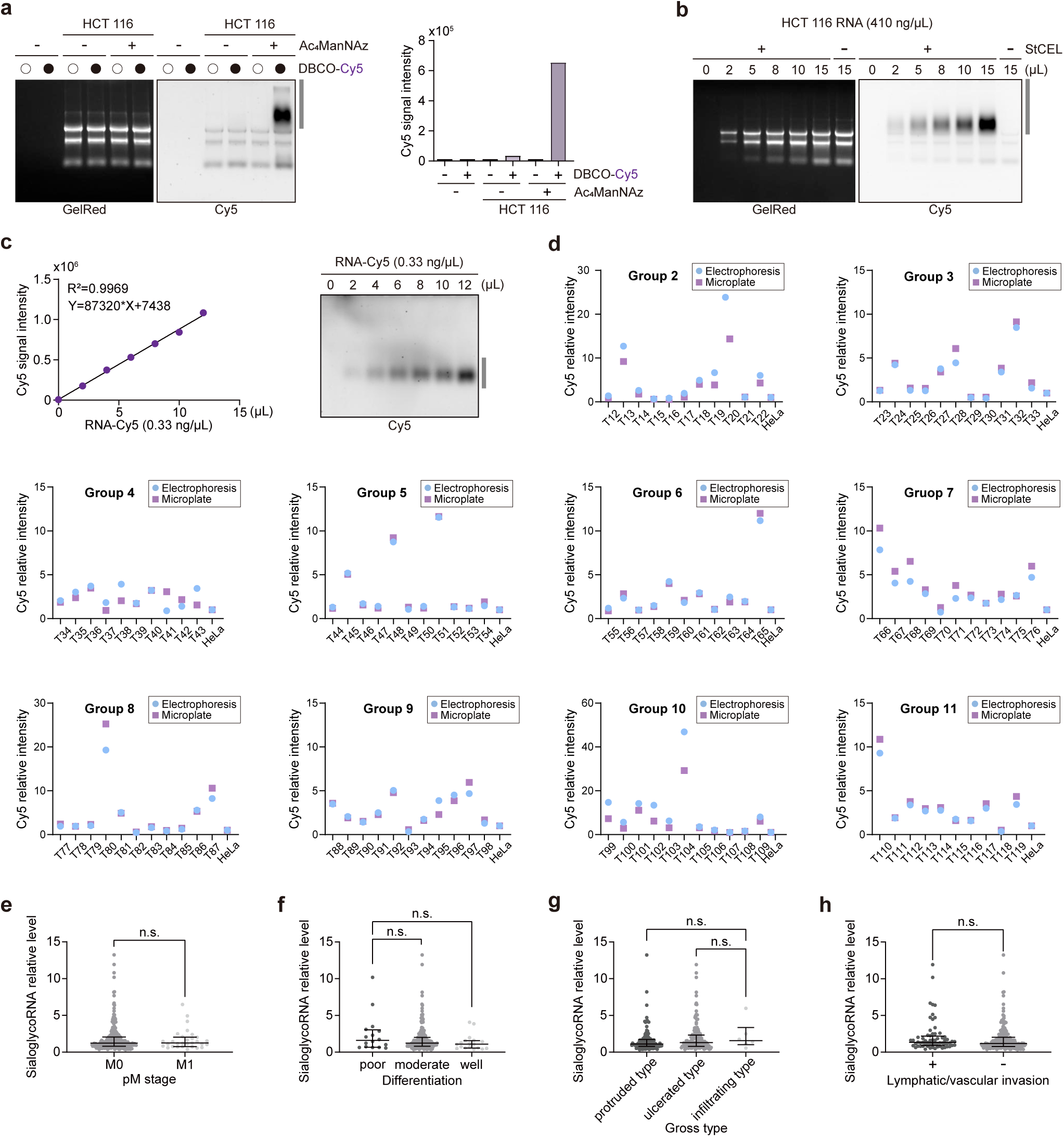
Detection of sialoglycoRNA in a large cohort of rectal tumor samples and analysis of clinicopathological correlations. **a**, Validation of DBCO-Cy5 conjugation efficiency following Ac_4_ManNAz labeling using gel electrophoresis (left) and microplate detection (right). **b**, RNA fluorescence imaging of StCEL-labeled HCT 116 total RNA at serial dilutions, captured with a multifunctional imager following gel electrophoresis. **c**, Linear correlation (left) between the signal values detected by the microplate reader and the amount of RNA-Cy5. Gel electrophoresis of the same sample was performed (right). **d**, Concordance validation between microplate reader and gel electrophoresis across 10 sample groups (Group 2-11; representative 108 of 256 samples shown). Corresponding tissue blot results are shown in **Supplementary Fig. 1e**. “HeLa” refers to the total RNA extracted from HeLa cells, which was used to correct the labeling efficiency of different groups. **e**, SialoglycoRNA levels stratified by pM stages. Statistical comparisons by Mann-Whitney test, *P*=0.9168. n.s. means not significant. **f**, SialoglycoRNA levels across differentiation (poor/moderate / well). Statistical comparisons by Kruskal-Wallis test, *P*=0.1750. n.s. means not significant. **g**, SialoglycoRNA levels across gross types (protruded/ulcerated/infiltrating). Statistical comparisons by Kruskal-Wallis test, *P*=0.1239. n.s. means not significant. **h**, SialoglycoRNA levels stratified by lymphatic/vascular invasion. Statistical comparisons by Mann-Whitney test, *P*=0.1206. n.s. means not significant.

**Extended Data Fig. 5.**
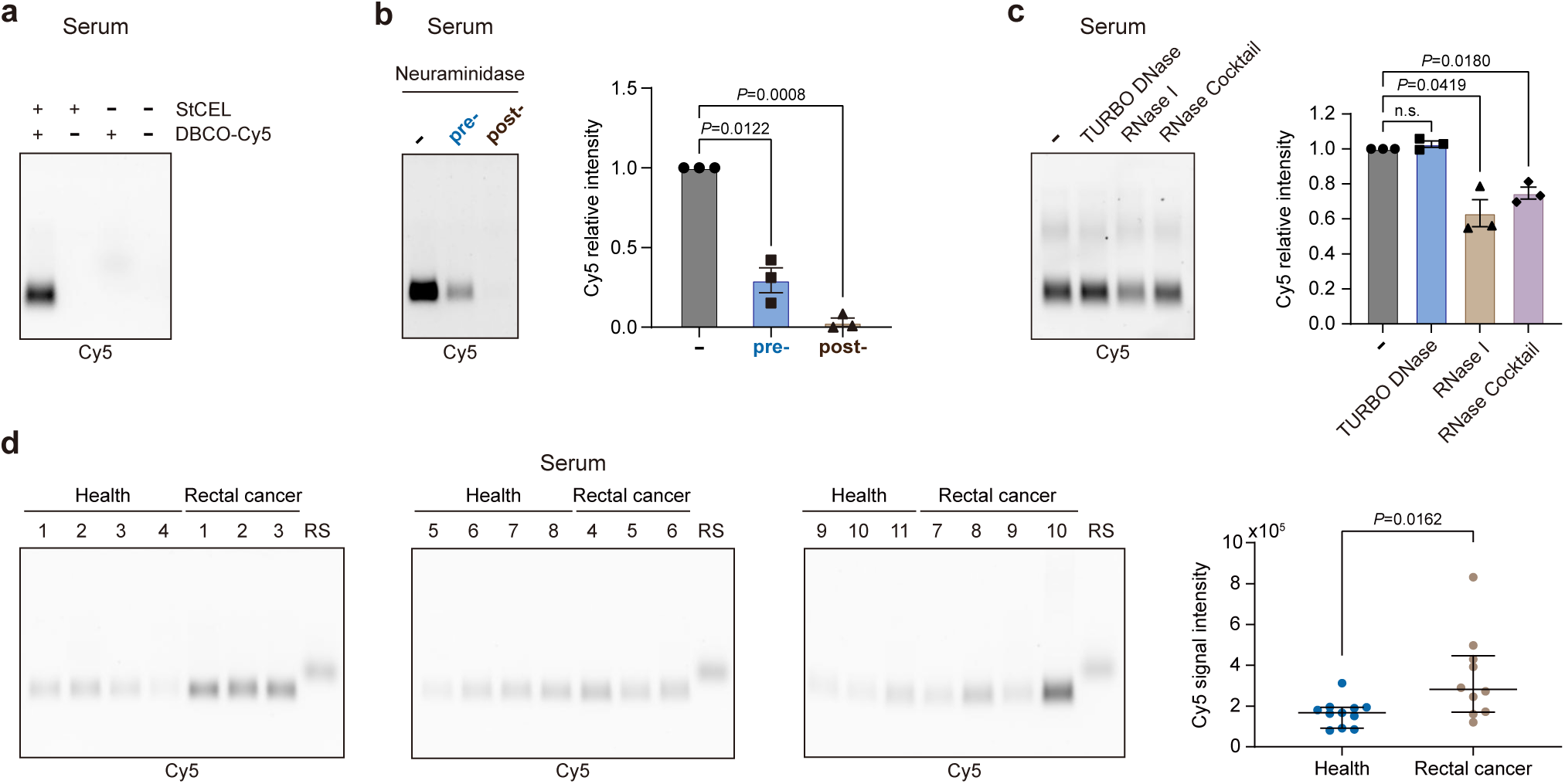
Detection of sialoglycoRNA in serum using StCEL. **a**, RNA fluorescent imaging of StCEL-labeled serum sialoglycoRNA. **b**, RNA fluorescent imaging of StCEL-labeled serum RNA from patients with rectal cancer treated with neuraminidase (α2-3,6,8,9 Neuraminidase A, left). Each datapoint (biological triplicate) was displayed with the mean ±SEM (right), and statistical analysis was performed with paired *t* tests. **c**, RNA blotting of StCEL-labeled serum RNA from patients with rectal cancer treated in vitro with RNase I, RNase cocktail (A/T1), TURBO DNase, or proteinase K (left). Each datapoint (biological triplicate) was displayed with the mean ±SEM (right), and statistical analysis was performed with paired *t* tests. **d**, Comparative serum sialoglycoRNA detection in healthy controls (n=11) versus rectal cancer patients (n=10). Left panel: in-gel fluorescent imaging. Right panel: densitometric quantification. Statistical analyses used an unpaired *t*-test. “RS” refers to the synthetic RNA standard (RNA-Cy5).

**Extended Data Fig. 6.**
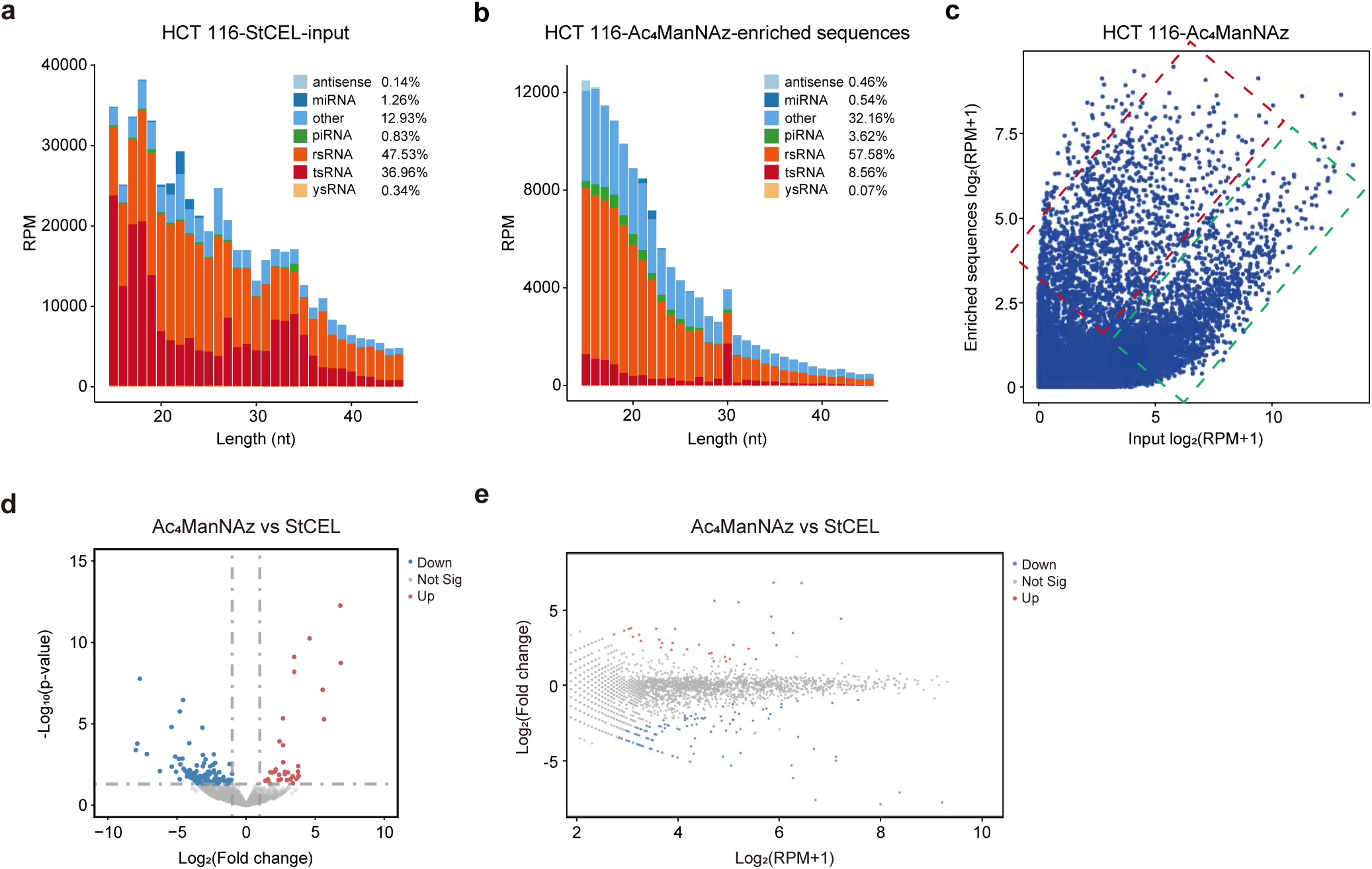
**Comparative analysis of sialoglycoRNA enriched by StCEL versus Ac_4_ManNAz from HCT 116 cells. a,b**, Length distribution of StCEL-input small RNA (**a**) and Ac_4_ManNAz-enriched sialoglycoRNA (**b**) from HCT 116 cells. **c**, Scatter plot comparing RNA sequence abundance in Ac_4_ManNAz-enriched sialoglycoRNA versus input small RNA. Red dashed boxes denote sequences enriched in purified sialoglycoRNA samples with positive abundance correlation, while green boxes indicate depleted sequences. **d**, Volcano plot depicting differentially enriched sialoglycoRNAs between Ac_4_ManNAz-and StCEL-enriched in HCT 116. **e**, Minus-versus-Add plot illustrating expression-level-dependent differences between Ac_4_ManNAz-and StCEL-enriched sialoglycoRNAs in HCT 116.

**Extended Data Fig. 7.**
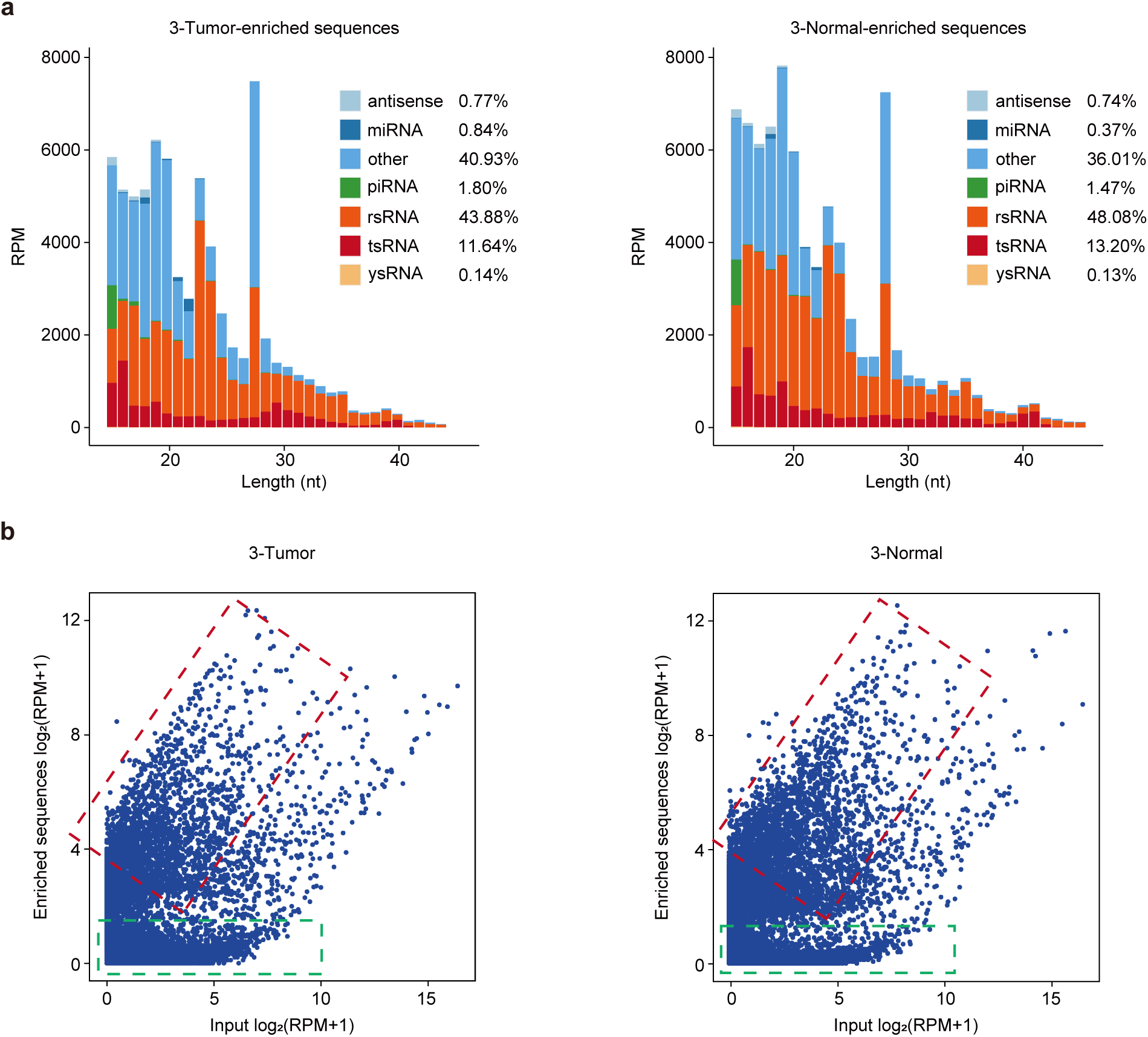
Comparative analysis of sialoglycoRNA from rectal tumor versus adjacent normal tissues by StCEL-based sialoglycoRNA-seq. a, Length distribution of StCEL-pull down sialoglycoRNAs in representative sample 3 with a pair of rectal tumor (left) *vs.* adjacent normal tissues (right). **b**, Scatter plots for StCEL-pull down sialoglycoRNA in representative sample 3 with pairs of tumor (left) and normal tissues (right).

**Extended Data Fig. 8.**
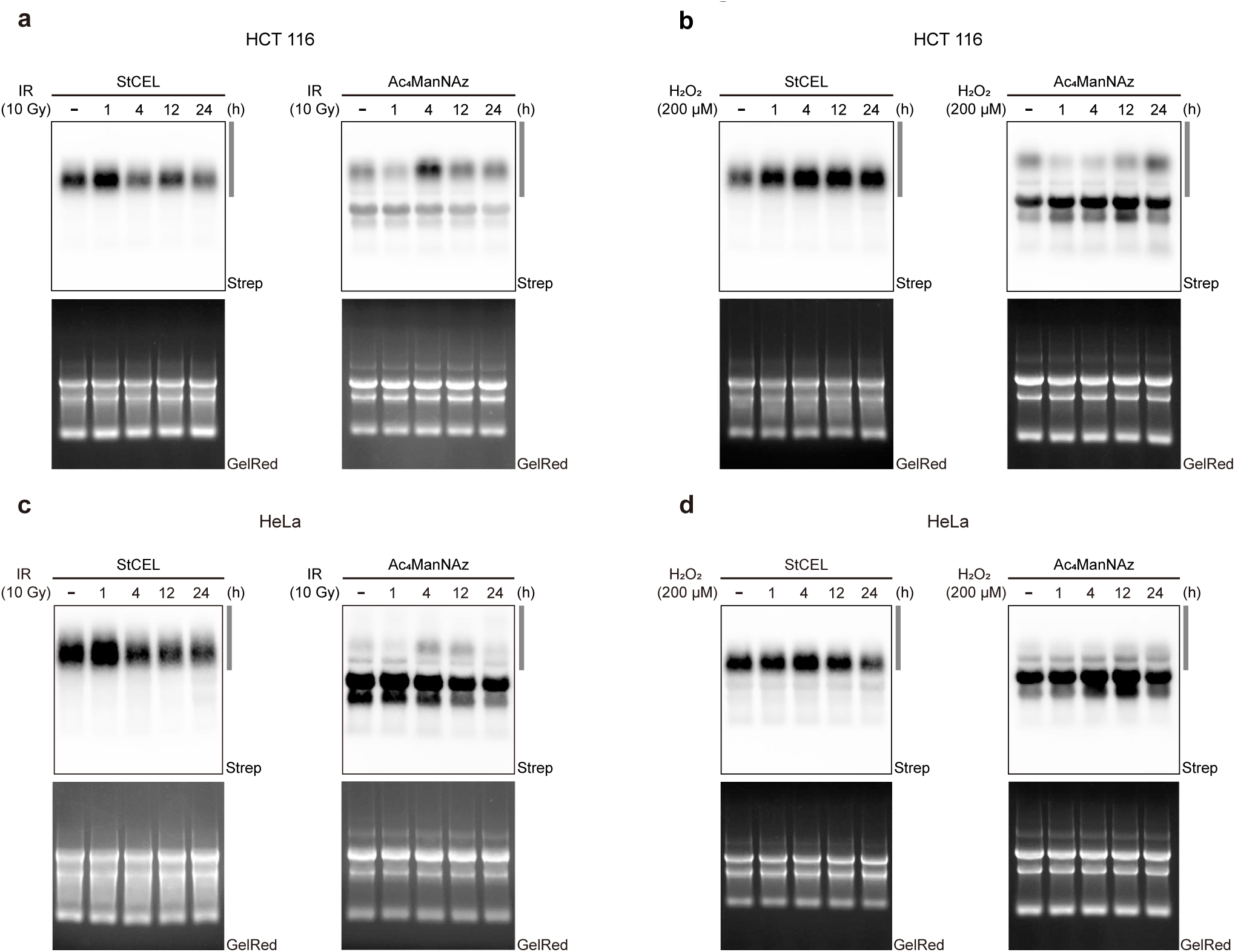
Detection of sialoglycoRNA dynamics in IR-or H₂O₂-treated cells using StCEL and Ac_4_ManNAz. **a,b**, RNA blotting of total RNA from HCT 116 cells collected at different time points after 10 Gy ionizing radiation **(a)** or 200 μM H_2_O_2_ **(b)** treatment, labeled with StCEL or Ac_4_ManNAz. **c,d**, RNA blotting of total RNA from HeLa cells collected at different time points after 10 Gy ionizing radiation **(c)** or 200 μM H_2_O_2_ **(d)** treatment, labeled with StCEL or Ac_4_ManNAz.

**Supplementary Fig. 1.**
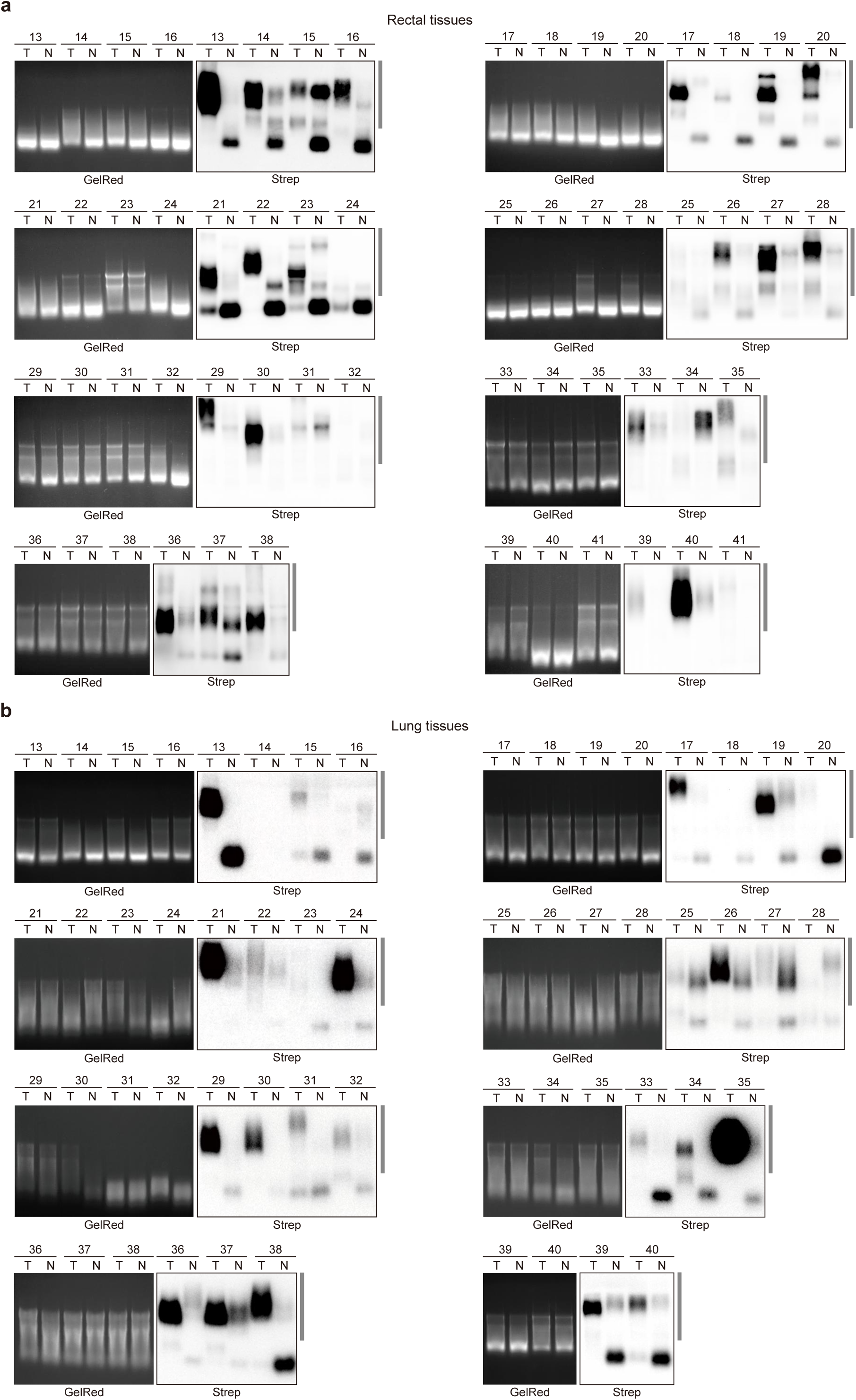
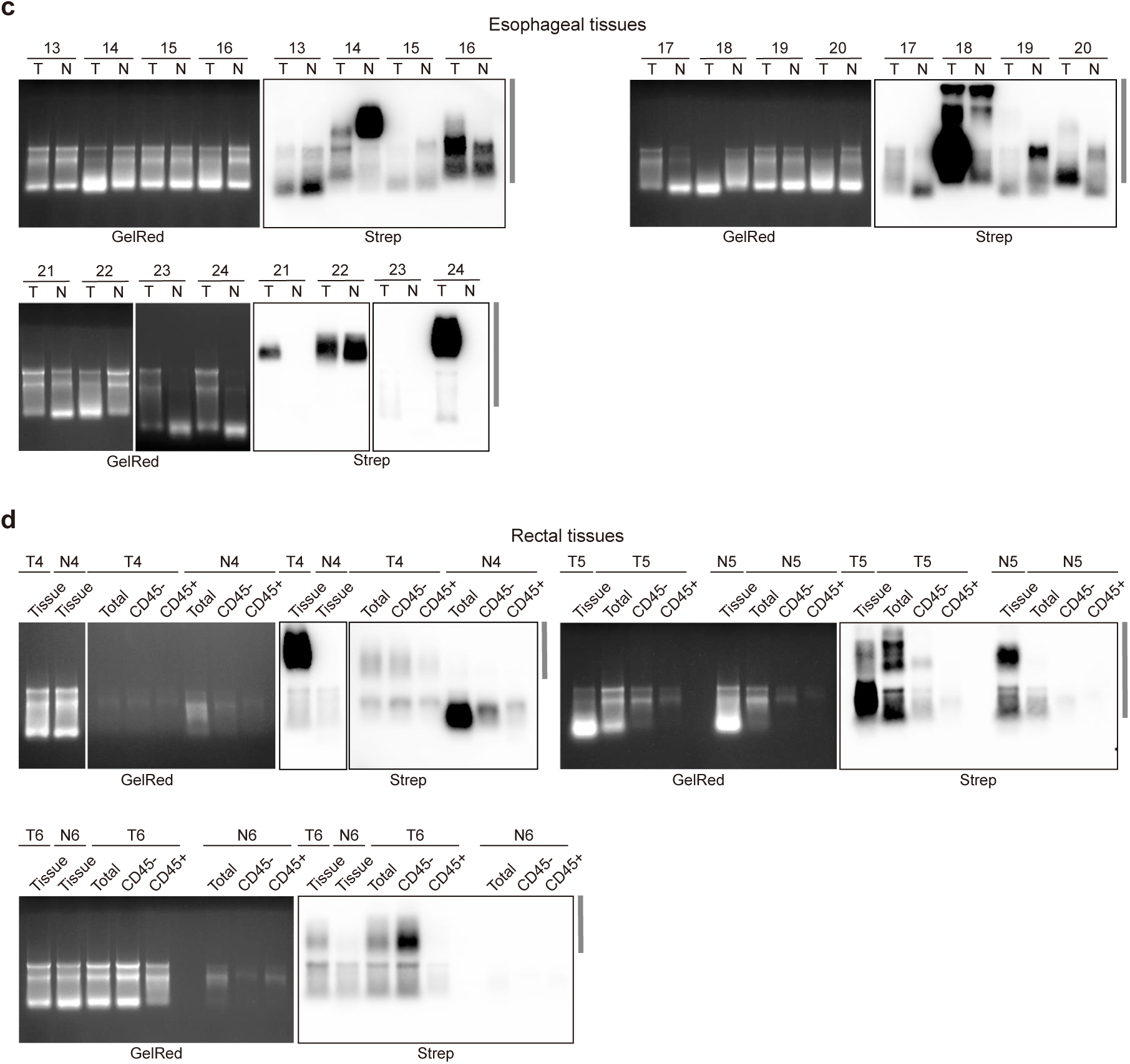
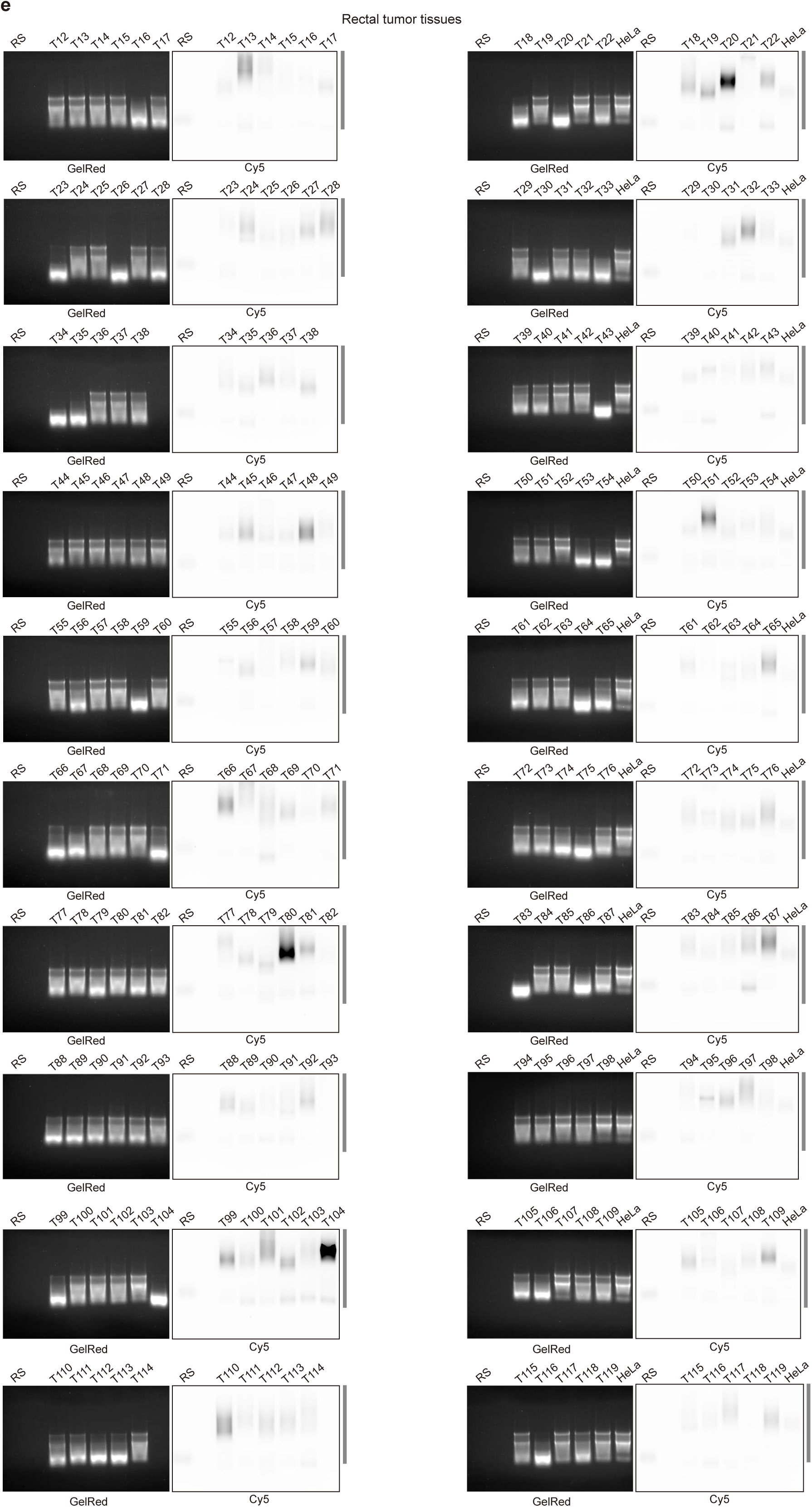
Data of sialoglycoRNA detection in human tissues. a, RNA blotting of the remaining samples in Fig. 3b **and Extended Data Fig. 2d** (29 of 41 pairs shown). **b,** RNA blotting of the remaining samples in Fig. 3c **and Extended Data Fig. 2e** (28 of 40 pairs shown). **c,** RNA blotting of the remaining samples in Fig. 3d **and Extended Data Fig. 2f** (12 of 24 pairs shown). **d,** RNA blotting of the remaining samples in Fig. 3f **and Extended Data Fig. 2h** (3 of 6 pairs shown). **e,** RNA blotting of the samples in **Extended Data Fig. 4d** (representative 108 of 256 pairs shown). “RS” referred to the synthetic RNA standard (RNA-Cy5). “HeLa” refers to the total RNA extracted from HeLa cells, which was used to correct the labeling efficiency of different groups.

